# Autophagic enhancer rescues Tau accumulation in a stem cell model of frontotemporal dementia

**DOI:** 10.1101/2024.09.30.615921

**Authors:** Farzaneh S. Mirfakhar, Jacob A. Marsh, Miguel A. Minaya, Stephen C. Pak, Gary A. Silverman, David H. Perlmutter, Shannon L. Macauley, Celeste M. Karch

## Abstract

Tau degradation is disrupted in neurodegenerative tauopathies, such as frontotemporal dementia (FTD), which may contribute to Tau aggregation. The prevailing hypothesis has been that Tau degradation is stymied due to an imbalance in proteostasis that occurs with age. Here, we used Airyscan super resolution imaging to illustrate that a pathogenic FTD mutation in the *MAPT* gene, which encodes Tau, is sufficient to alter multiple steps of the autophagy lysosomal pathway and impair Tau degradation. We discovered lysosomes clogged with both Tau and phosphorylated Tau, stalled lysosome motility, disrupted molecular motors, enhanced autophagic flux, and slowed cargo degradation in mutant Tau neurons. Treatment of mutant Tau neurons with a small molecule autophagy enhancer drug increases autophagic flux and cargo degradation, reduces phospho-Tau levels, and reduces Tau accumulation in lysosomes without restoring defects in lysosomal motility. This study reveals novel effects of mutant Tau and provides a window through which therapeutic treatments targeting autophagy may promote Tau homeostasis.

## Introduction

The dysregulation of protein degradation pathways has been implicated across neurodegenerative disorders ^1, 2, 3^. In the context of frontotemporal lobar dementia with Tau inclusions (FTLD-Tau), pathological protein aggregates are composed of hyperphosphorylated microtubule-associated protein Tau ^4, 5^. The Tau protein, encoded by the *MAPT* gene, serves to stabilize and organize microtubules within neurons ^6, 7^. Mutations within the *MAPT* gene cause familial forms of FTLD-Tau, leading to altered Tau protein structure ^8, 9, 10^ and Tau aggregation ^5,10^. However, the mechanism by which these events occur remains uncertain.

As polarized, terminally differentiated cells, neurons encounter distinctive challenges in maintaining the quality and composition of their proteome ^11^. Unlike proliferating cells, neurons cannot dilute proteotoxins through cell division ^12^. Consequently, neurons tightly regulate proteostasis through parallel mechanisms including the proteosome, macroautophagy, chaperone mediated autophagy, and endo-microautophagy ^1, 13, 14, 15^. Defects in neuronal autophagy have been implicated in the pathogenesis of numerous neurodegenerative disorders ^16^. However, whether disease-causing mutations are sufficient to disrupt these pathways in the absence of robust protein aggregates remains to be understood.

In autophagy, autophagosomes engage with dynein for retrograde transport to the soma, where they undergo maturation into degradative compartments through fusion with LAMP1-positive lysosomes, ultimately leading to cargo degradation ^1, 15^. Numerous motor-associated proteins have been identified to regulate the dynein-driven retrograde transport of autophagic vesicles along neurites ^17, 18, 19, 20, 21^. Alterations in these motor-associated proteins can disrupt the transport of autophagic vesicles and impede their maturation into degradative lysosomes ^22^. Consequently, there exists a link between lysosome localization and motility, with its degradative capacity and function ^18, 20, 23^. Therefore, defects in lysosomal trafficking pathways can perturb neuronal clearance mechanisms, thereby promoting Tau aggregation and contributing to neurodegeneration ^3^.

Here, we sought to determine whether alternations in the autophagy lysosomal pathway are early features in neurons from *MAPT* mutation carriers and whether restoring these pathways is sufficient to rescue Tau dysfunction. We found that *MAPT* p.R406W is sufficient to alter multiple steps of the autophagy lysosomal pathway and impair Tau degradation. We discovered that mutant Tau neurons have Tau and pTau-laden lysosomes, stalled lysosome motility, disrupted molecular motors, enhanced autophagic flux, and slowed cargo degradation. Treatment of mutant Tau neurons with an autophagy enhancer drug G2 series increases autophagic flux and cargo degradation, reduces pTau levels, and reduces Tau accumulation in lysosomes without restoring defects in lysosomal motility. Together, our findings point to novel effects of mutant Tau that can be rescued by targeting autophagy.

## Results

### Post translational modifications impact Tau localization in lysosomes in human neurons

Post-translational modifications (PTMs) on Tau, particularly phosphorylation, play an important role in the normal function of Tau to bind to and stabilize microtubules ^24, 25^. Hyperphosphorylation of Tau alters its conformation, promoting Tau aggregation ^26, 27, 28^. Phosphorylated Tau may be sequestered into autophagic vacuoles and targeted for degradation through the autophagy-lysosome pathway ^13, 29^. The intricate interplay between Tau phosphorylation and degradation pathways highlights the significance of PTMs in regulating Tau homeostasis ^13^. While Tau processing in degradative vesicles has been described, these studies have relied on overexpression systems. Little is known of how human neurons process endogenous Tau and how endogenous Tau PTMs impact these degradative pathways. To understand the impact of Tau phosphorylation on Tau degradation in lysosomes, we used patient-derived iPSC-derived neurons harboring a doxycycline-inducible neurogenin-2 (NGN2) cassette stably integrated in the AAVS1 locus (**Supplemental Figure 1-2**). NGN2 expression in iPSCs results in a homogeneous population of excitatory neurons within seven days ^30^. These neurons expressed the neuronal markers Microtubule-Associated Protein 2 (MAP2) and β-tubulin III (Tuj1) by DIV7 and expressed synapsin by DIV14 (**Supplemental Figure 3**).

To understand where endogenous Tau localizes within the endolysosomal compartments of human neurons, we visualized total Tau (Tau5) and phosphorylated Tau (AT180; pThr231) in LAMP1-positive (LAMP1+) vesicles of DIV14 neurons using immunocytochemistry and Airyscan super resolution microscopy (**Figure 1A**). LAMP1+ vesicles were positive for cathepsin D and appeared as donut-shaped structures (**Figure 1A**; **Supplemental Figure 4**); thus, these LAMP1+ structures are likely lysosomes. Super resolution images of LAMP1+ vesicles showed that Tau and pTau localize to different compartments in the lysosome (**Figure 1A-B**). Signal intensity measurements illustrate that Tau localization was enriched within the lysosomal lumen (**Figure 1B-C**), while pTau localization was enriched at the lysosomal membrane (**Figure 1B and 1D**). We discovered a difference in the retention of Tau and pTau in lysosomes, which may point to a difference in degradative efficiency: 70% of lysosomes were free of Tau (Tau- or empty) while only 40.7% of lysosomes were free of pTau (**Figure 1B and 1E**). Thus, our findings point to differences in the way wild-type neurons handle Tau and pTau in the lysosome, which may ultimately lead to differences in degradation efficiency (**Figure 1F**).

**Figure 1:**
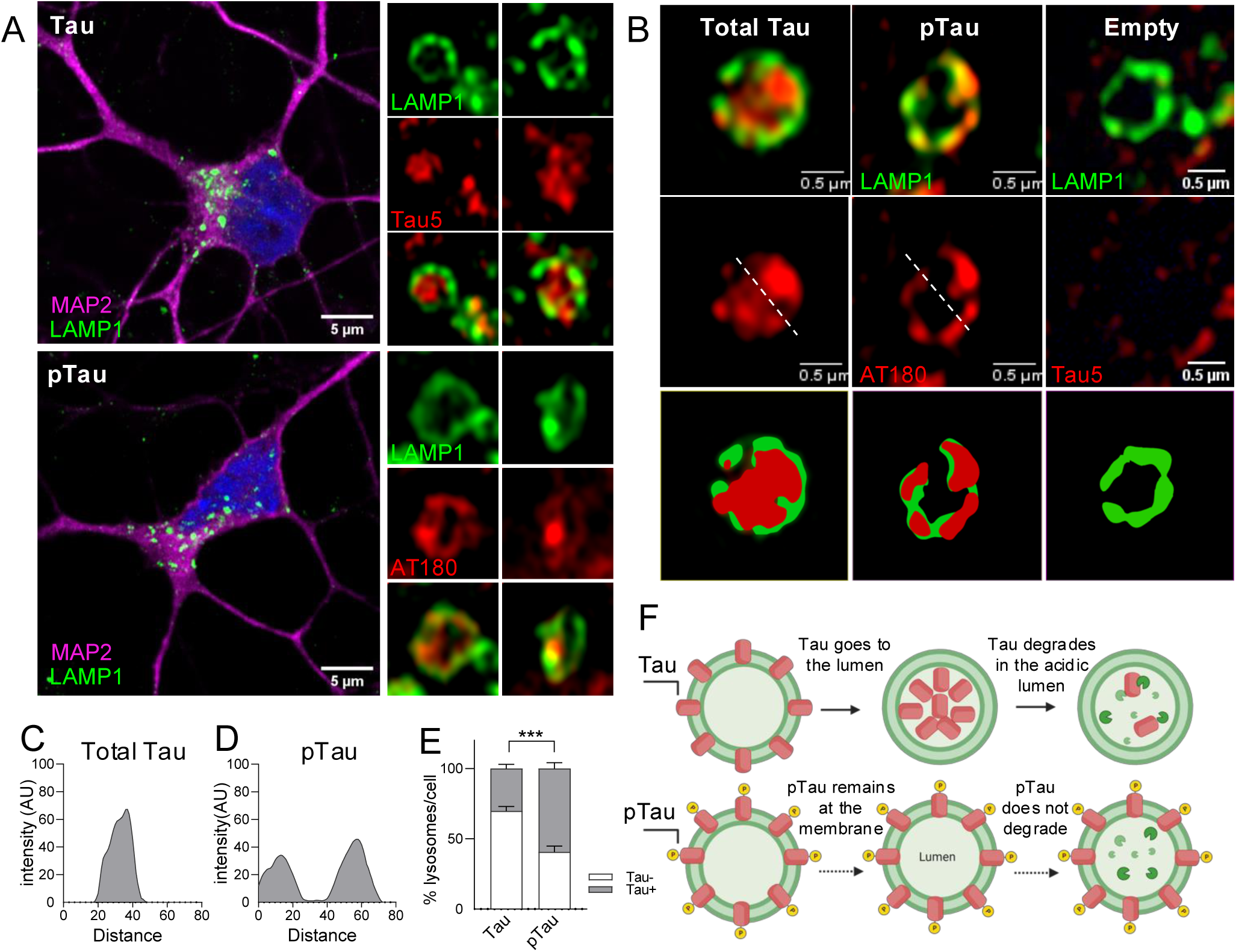
Tau phosphorylation shifts lysosomal localization of Tau in human neurons. A. iPSC-derived NGN2 neurons (control) were cultured until DIV14, immunostained for LAMP1+ vesicles (green), MAP2 (magenta), Tau (Tau5; red) and pTau (AT180; red), and imaged by Airyscan confocal microscopy. Scale bar 5µm. Right inset, LAMP1+ vesicles co-stained for total Tau and pTau AT180. Scale bar 0.5µm. B. Tau and pTau localization within LAMP1+ vesicles differ. Dashed line used as reference for subsequent quantification. LAMP1+ vesicles (green) co-stained for total Tau (Tau5; red) and pTau (AT180; red). Scale bar 0.5µm. C-D. Quantification of fluorescent intensity of Tau (C) and pTau (AT180; D) in LAMP1+ vesicles across the dashed line in B. Quantification based on 3 independent experiments. E. Quantification of the percentage of LAMP1+ vesicles per cell that contain total Tau or pTau (Tau+ or free of Tau, Tau-). Graph represents mean ± SEM (186 vesicles quantified from 3 independent experiments). Two-tailed Mann-Whitney *U* test. **, p=0.0043. F. Diagram of Tau and pTau localization in lysosomes based on our findings.

### Tau degradation is stalled in MAPT p.R406W neurons

Given the finding that Tau and pTau differ in their lysosomal localization in wild-type neurons, we sought to understand the impact of FTD-causing *MAPT* mutations on Tau phosphorylation and Tau degradation in lysosomes. Studying Tau degradation in neurons has been challenging due to the lack of proper human cell models, resulting in the over reliance of Tau overexpression in immortalized cell lines^31, 32^. NGN2 was engineered into the AAVS locus of an iPSC line from a patient carrying the FTD-causing *MAPT* p.R406W mutation and its isogenic, WT pair (**Figure 2A; Supplemental Figure 1-2**). *MAPT* p.R406W neurons (DIV14) produced significantly more Tau phosphorylated at pThr231 (AT180; **Figure 2B-C**) and pSer202/pThr205 (AT8; **Figure 2D-E**) compared to isogenic control neurons.

**Figure 2:**
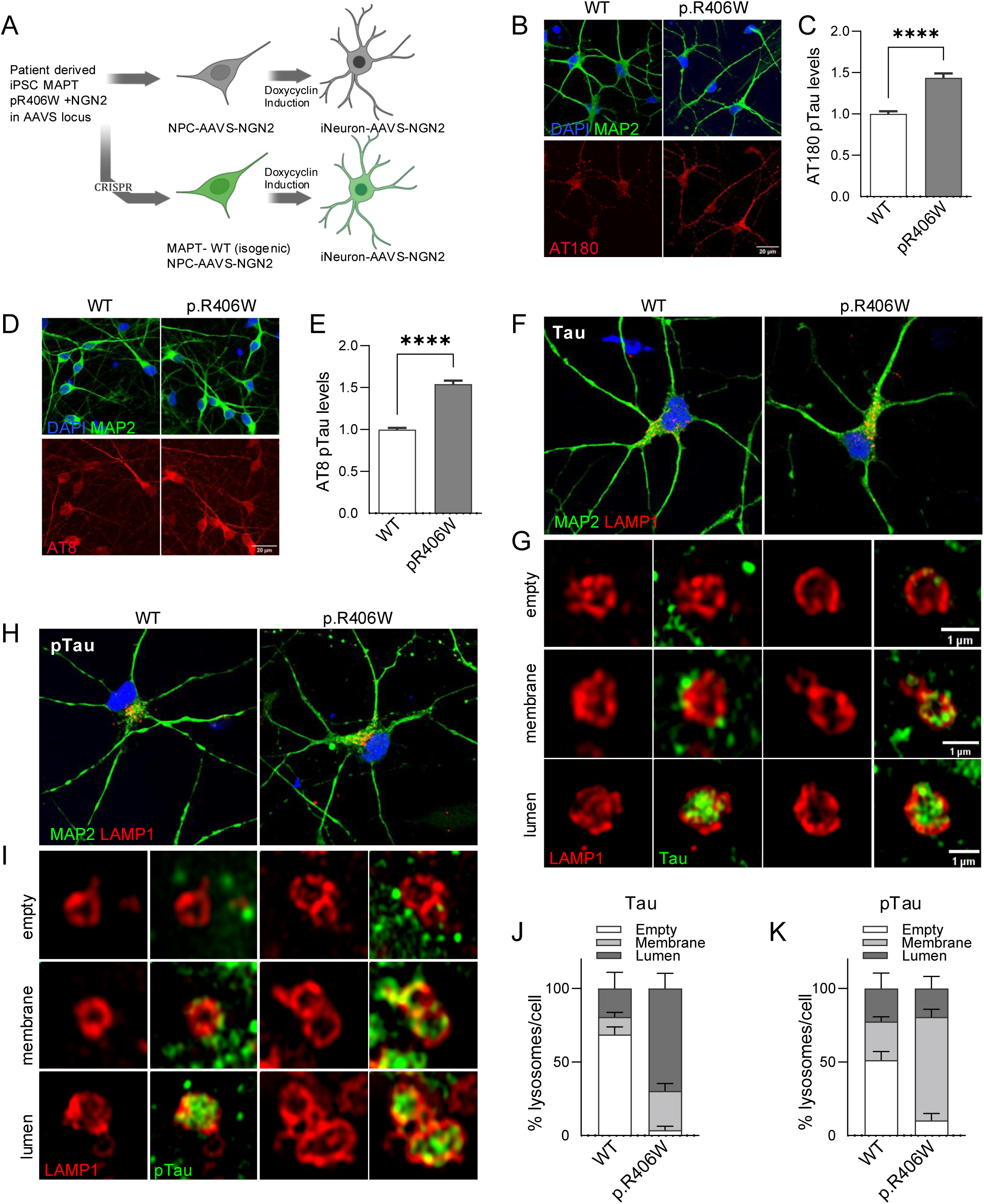
Tau accumulates in lysosomes in *MAPT* p.R406W neurons. A. Diagram of stem cell model used in this study. Human derived iPSC with NGN2 integrated in the AAVS locus were differentiated into neurons. *MAPT* mutation carrier (p.R406W) and CRISPR/Cas9-corrected isogenic control (wild-type; WT) were differentiated into cortical neurons upon doxycycline induction. B. p.R406W neurons display more Tau phosphorylated at Thr231. Neurons were defined with MAP2 staining (green). WT and p.R406W neurons were stained for pTau (AT180; red). Scale bar 20µm. C. Quantification of AT180 pTau relative mean fluorescence intensity. Data are represented as mean ± SEM. Total 159 cell quantified from 3 independent experiments. Two-tailed Mann-Whitney *U* test, ****, p<0.0001. D. p.R406W neurons display more Tau phosphorylated at Ser202/Thr205. Neurons were defined with MAP2 staining (green). WT and p.R406W neurons stained for pTau (AT8; red). Scale bar 20µm. E. Quantification of AT8 pTau relative mean fluorescence intensity. Data are represented as mean ± SEM. Total 418 cells quantified from 4 independent experiments. Two-tailed Mann-Whitney *U* test. ****, p<0.0001. F. WT and p.R406W were stained for LAMP1 (red) and MAP2 (green). G. Magnified images of Lamp1+ vesicles (red) and total Tau (Tau5; green). Scale bar 1µm. Representative images of LAMP1+ vesicles empty of total Tau (top panel), total Tau staining in the lysosomal membrane (middle panel), and total Tau staining in the lysosomal lumen (bottom panel). H. WT and p.R406W were stained for LAMP1 (red) and MAP2 (green). I. Magnified images of LAMP1+ vesicles (red) and pTau (AT180; green). Representative images of LAMP1+ vesicles empty of pTau (top panel), pTau staining in the lysosomal membrane (middle panel), and pTau staining in the lysosomal lumen (bottom panel). J-K. Quantification of the percentage of LAMP1+ vesicles per cell. Data are mean ± SEM. Total 186 vesicles quantified from 3 independent experiments. Two-way ANOVA mixed model, WT:Empty vs. p.R406W:Empty **, p=0.0010, WT:Lumen vs. p.R406W:Lumen ***, p= 0.0001, WT:Empty vs. WT:Membrane **, p= 0.0040, WT:Empty vs. WT:Membrane *, p=0.0192. p.R406W: Empty vs. p.R406W: Membrane *, p= 0.0169, p.R406W:Membrane vs. p.R406W:Lumen *, p= 0.0159. J. LAMP1+ vesicles empty of total Tau or with total Tau in the membrane vs lumen. K. LAMP1+ vesicles empty of pTau or with pTau in the membrane vs lumen.

To determine whether *MAPT* p.R406W alters Tau localization in lysosomes, we immunostained *MAPT* p.R406W neurons and isogenic controls for Tau (Tau5) and pTau (AT180) at DIV14. Tau was enriched in the lysosomal lumen in *MAPT* p.R406W and its isogenic control (**Figure 2F-G**); however, mutant neurons exhibited significantly more Tau (p.R406W:69.6% vs. WT:17.64%) in the lumen than the isogenic control neurons (**Figure 2F-G, 2J**). We also observed a modest increase in membrane bound Tau in the mutant neurons compared to the isogenic controls (p.R406W: 26.7% vs. WT: 11.8%; **Figure 2F-G, 2J**). *MAPT* p.R406W neurons exhibited significantly more pTau in the lysosomal membrane than isogenic controls (p.R406W: 70.3% vs. WT: 26.2%; **Figure 2H-I, 2K**). Strikingly, *MAPT* p.R406W neurons had fewer lysosomes free of Tau (p.R406W: 3.6% vs. WT: 68.7%) and pTau (p.R406W: 10.3% vs. WT: 51.2%) compared to isogenic controls. Together, our findings suggest that degradation of Tau and pTau in lysosomes is stalled in mutant neurons.

### Endolysosomal pathways are disrupted in MAPT p.R406W neurons

To begin to understand the mechanisms underlying Tau accumulation in lysosomes in *MAPT* p.R406W neurons, we performed transcriptomic analyses in mutant neurons and isogenic controls ^33^. Genes enriched in synaptic function and endolysosomal function were among the most significantly differentially expressed genes ^33^. In *MAPT* p.R406W neurons, endolysosomal genes were enriched in pathways associated with phagosome maturation, regulation of autophagy, autophagosome, and vesicle transport along the microtubule (**Figure 3A**; **Supplemental Figure 5**). Autophagosome clearance is a multistep process involving vesicular trafficking, fusion, and degradation ^34^. Autophagy impairment can lead to a disruption in normal fusion and degradation processes within the cell ^35, 36, 37^.

**Figure 3:**
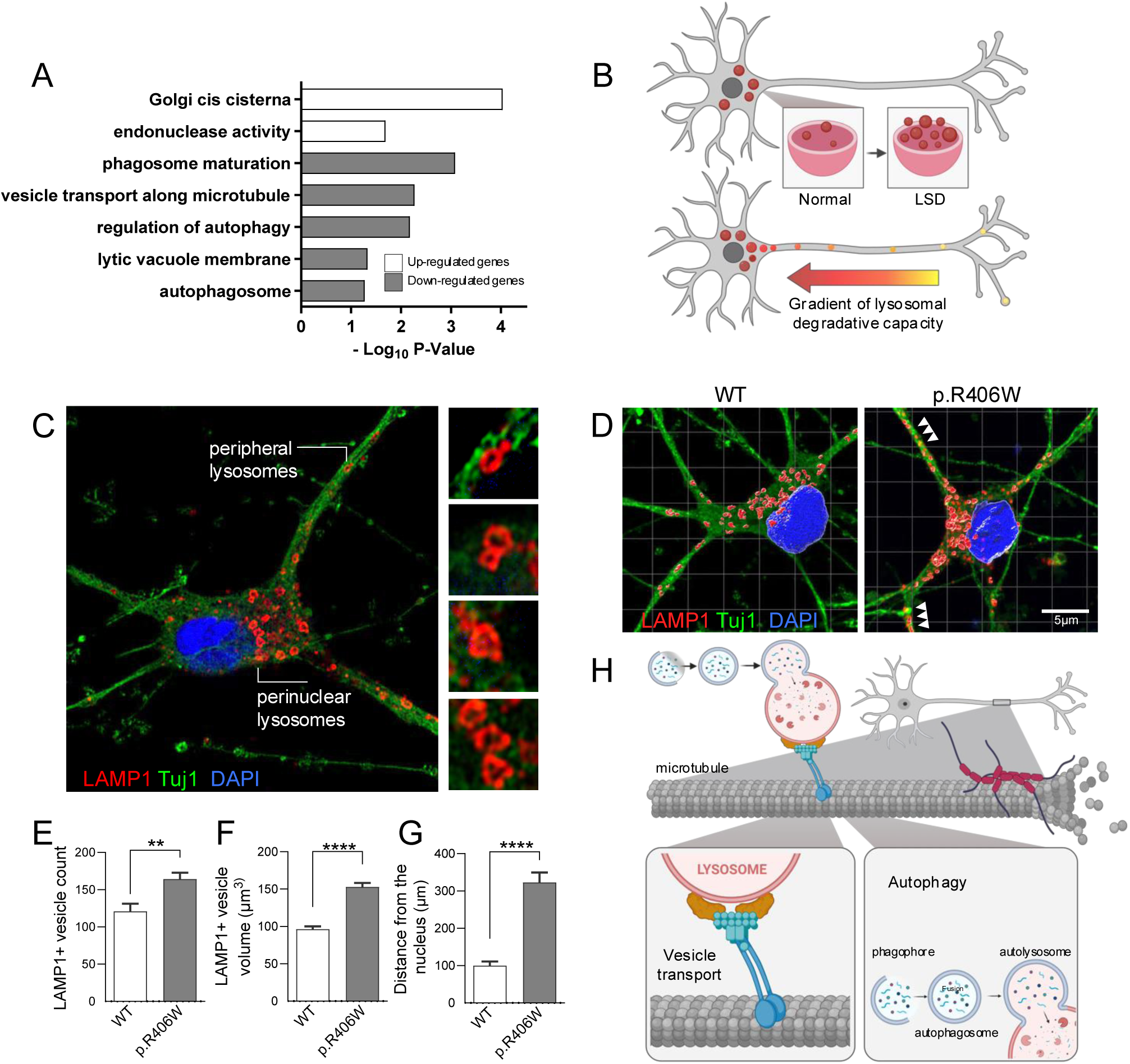
Endolysosomal pathways are disrupted in *MAPT* p.R406W neurons. A. RNA sequencing analysis of *MAPT* p.R406W neurons compared to isogenic controls revealed differentially expressed genes enriched in GO term associated with the endolysosomal pathway. White bars, pathways with genes significantly upregulated (FDR-BH ≤ 0.05) in *MAPT* p.R406W neurons. Gray bars, pathways with genes significantly downregulated (FDR-BH ≤ 0.05) in *MAPT* p.R406W neurons. B. Schematic of lysosomal size and localization relative to predicted degradative capacity. C. Representative Tuj1+ (green) neurons at DIV14, stained for LAMP1 (red) for detecting peripheral lysosomes vs perinuclear lysosomes. Right, magnified images of LAMP1+ donut-shaped structures (red) distributed in neurons. D. Tuj1+ (green) neurons at DIV14. *MAPT* p.R406W or isogenic controls (WT) were stained for LAMP1 (red). 3D reconstructed of Airyscan microscopy images was performed by Imaris. White arrows indicate lysosomes located at an extended distance from the soma. Scale bar, 5µm. E-G. Lysosomal morphology quantified in *MAPT* p.R406W neurons compared with isogenic controls (WT). Data are mean ± SEM from 3 independent experiments. Two-tailed Mann-Whitney *U* test. E. Quantifications of lysosome density (measurement of total number of LAMP1+ puncta per Tuj1+ cells, quantified by Imaris.19 cells per genotype, **, p=0.0048). F. Lysosome volume (measurement of lysosome volume from z-stack images, quantified by Imaris, number of lysosomes quantified; *MAPT* WT n=208; *MAPT* p.R406W n=318; ****, p<0.0001. G. Lysosome distance from the nucleus (measurement of shortest distance of LAMP1+ puncta to DAPI+, number of lysosomes quantified: *MAPT* WT n=164; *MAPT* p.R406W n=214, ****, p<0.0001). H. Diagram of hypothesized disrupted pathways.

When autophagy is impaired, undegraded material may accumulate within the lysosomes, causing them to enlarge ^38, 39^. Lysosomal enlargement is an indication of undigested material reminiscent of the pathophysiology observed in lysosomal storage disorders (LSD; **Figure 3B**) ^40, 41^. To evaluate the impact of *MAPT* p.R406W on lysosomal size, we used Airyscan super-resolution microscopy to visualize LAMP1-positive, ring-shaped structures of different sizes at the cell periphery along the neurites as well as at the perinuclear region (**Figure 3C**). Quantification of LAMP1+ vesicle density and volume revealed increased lysosomal density (vesicle count) and volume in *MAPT* p.R406W neurons compared to the isogenic control neurons (**Figure 3D-F**). An increase in lysosomal volume is consistent with the accumulation of undigested material, including Tau and pTau, in *MAPT* p.R406W neurons.

Our transcriptomic analyses identified defects in vesicle transport along the microtubule which may influence lysosomal positioning within the neurons. Lysosomal distance from the nucleus has been shown to influence the degradative capacity of lysosomes (**Figure 3B-C**) ^42^. To gain insight into the impact of *MAPT* p.R406W on lysosomal positioning along the neurites, we measured the LAMP1+ vesicle distance from the nucleus. Lysosomes in *MAPT* p.R406W neurons were located further from the nucleus compared to the lysosomes in isogenic control neurons (**Figure 3D, 3G**). The peripheral localization of lysosomes in mutant neurons may reflect aberrant vesicle trafficking, suggesting a potential defect in retrograde trafficking (**Figure 3H**) ^21, 43^. Taken together, molecular and morphologic evidence suggest that *MAPT* p.R406W induces defects in autophagy and lysosomal trafficking in human neurons, which may impact degradative capacity.

### MAPT p.R406W neurons exhibit defects in lysosomal transport

Many lysosomal functions, including degradative capacity, are influenced by their positioning and motility ^18, 20^. Although some lysosomes are relatively static, others move along microtubules between the soma and the periphery of the cell ^18^. Our data suggest that lysosomes are farther away from the soma in *MAPT* p.R406W neurons (**Figure 3G**). Given the relationship between Tau and microtubules, we hypothesized that *MAPT* p.R406W disrupts lysosomal trafficking. Lysosomal motility was monitored using time-lapse confocal live-cell imaging of lysotracker-positive (lysotracker+) vesicles (**Figure 4A**). In *MAPT* p.R406W neurons, lysotracker+ vesicles traveled significantly shorter distances compared to isogenic controls (**Figure 4A-B**). Lysotracker+ vesicles also exhibited reduced velocity in mutant neurons compared to isogenic controls (**Figure 4A, 4C**). Together, these findings suggest that *MAPT* p.R406W impacts lysosomal motility.

**Figure 4:**
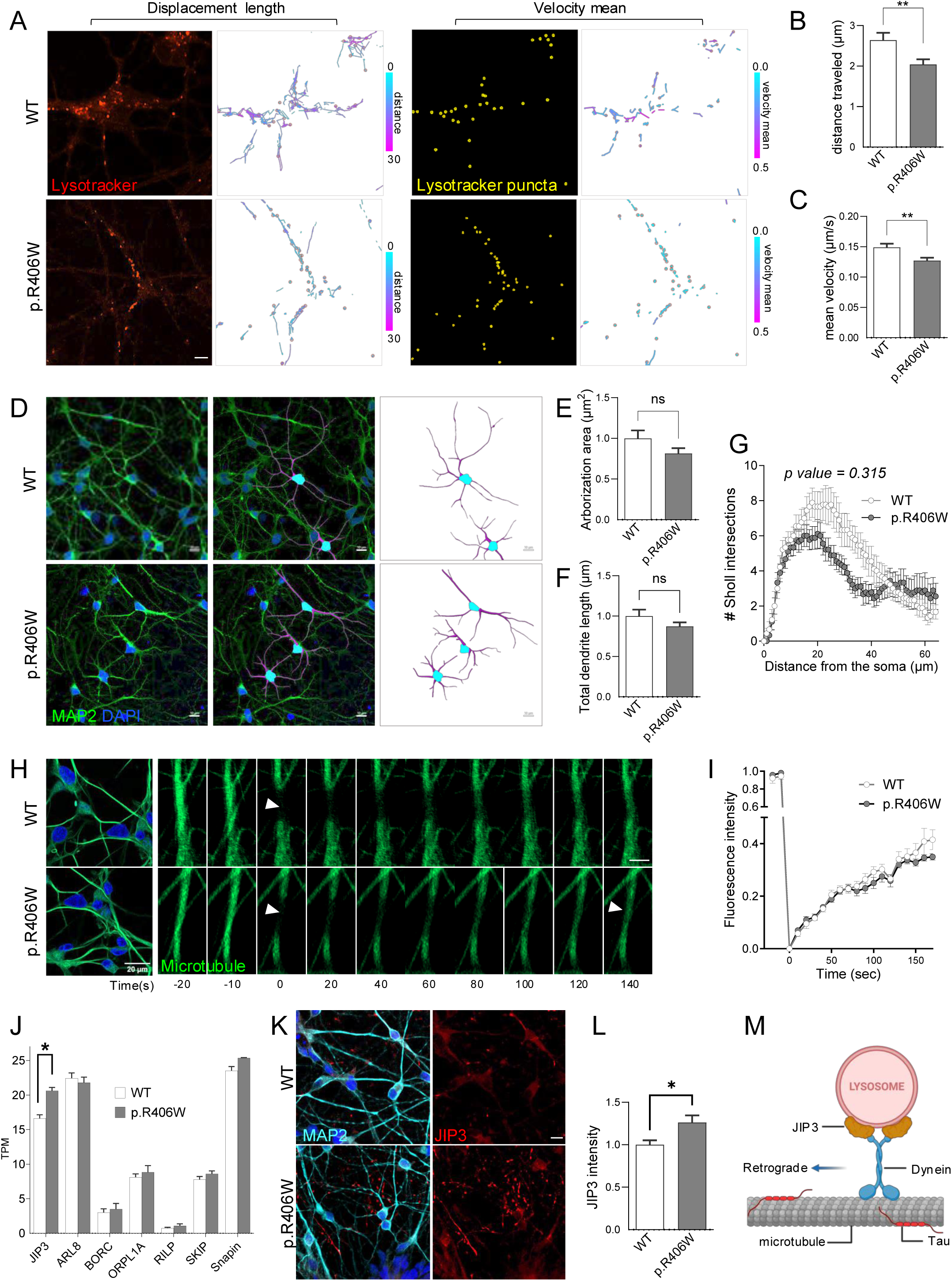
*MAPT* p.R406W neurons exhibit defects in lysosomal transport. *MAPT* p.R406W neurons compared with isogenic controls (WT) at DIV14. A. Displacement and velocity of acidic vesicles (identified with LysoTracker) was visualized using live imaging. Left: Lysotracker positive (red) acidic vesicles in *MAPT* WT and p.R406W were tracked for displacement length by Imaris (right, tracks are color-coded with cyan being shorter distances and magenta longer distances), Scale bar 10µm. Right: Lysotracker + puncta detected (yellow) and tracked by Imaris (right, tracks are color-coded with cyan being slower velocity and magenta faster velocity), Scale bar 10µm. B. Quantification of relative distance traveled by lysotracker+ vesicles in A. Data are mean ± SEM (*MAPT* WT n=510 and p.R406W n=856 lysotracker+ vesicles tracked from 3 independent experiments), t test, ***, *p=0.0048*. C. Quantification of relative mean velocity of lysotracker + vesicles in A. Data are mean ± SEM (*MAPT* WT n=336 and p.R406W n=415 lysotracker+ vesicles tracked from 3 independent experiments), t test, **, *p=0.0038*. D. Dendritic complexity was evaluated by capturing neuronal morphology of *MAPT* p.R406W and isogenic control neurons with MAP2 (green). 3D reconstruction by Imaris (magenta) based on MAP2 signal (green). Scale bar 10µm. E. Quantification of arborization area (p=0.195). F. Total dendrite length was quantified by measuring the total lengths of all branches per dendritic compartment in *MAPT* WT and p.R406W neurons (p=0.385). G. Sholl analysis. Quantification of the number of intersections at concentric circles of increasing radii centered around the soma every 10μm (p=0.315). E-G. Data are represented as mean ± SEM (*MAPT* WT n=30 and p.R406W n=35 cells were quantified). Two-tailed Mann-Whitney *U* test (E and F) and mixed model ANOVA (G). H. FRAP analysis of microtubules (Viafluor, green) in dendritic sections of *MAPT* p.R406W and isogenic control neurons, time-lapse sequences are shown to demonstrate the recovery of fluorescence in bleached ROI. I. Quantification of fluorescence recovery after photobleaching of microtubules signals in *MAPT* WT and p.R406W. The curves depict the average intensity at different time points normalized to the first frame (number of cells quantified for several ROIs in *MAPT* WT n=9 and pR406W n=8), mixed model ANOVA, p=0.22. J. Expression level of motor-associated genes detected by RNA-seq in *MAPT* p.R406W and isogenic control neurons. Expression levels are reported as transcripts per million (TPM) and have been log2-transformed and normalized across all the cell samples (p=0.023). K. *MAPT* p.R406W and isogenic control neurons were immunostained for molecular motor protein JIP3 (red). Scale bar 10µm. L. Quantification of JIP3 mean fluorescence intensity. Data are mean ± SEM (*MAPT* WT n=23 and p.R406W n=25 cells quantified from 3 independent experiments), Two-tailed Mann-Whitney *U* test, *p=0.0358. M. Schematic illustrating molecular motor and motor-associated proteins involved in lysosomal transport in neuronal processes ^49^.

To determine whether the observed alterations in lysosomal distribution and motility in *MAPT* p.R406W neurons were attributed to mutation-induced alterations in cell morphology, we quantified dendritic arborization, dendritic length and nodes in MAP2-immunostained neurons (**Figure 4D**). Neuronal arborization, dendritic length, and the number of nodes (measured by Sholl analysis) were similar between *MAPT* p.R406W and isogenic control neurons (**Figure 4D-G**). Thus, *MAPT* p.R406W likely does not impact lysosomal motility due to defects in neuronal morphology.

Vesicles move along microtubules and the Tau protein promotes microtubule stability. Therefore, we tested whether *MAPT* p.R406W alters microtubule stability, to account for the lysosomal trafficking defects. Microtubule dynamic turnover was measured in live cells by quantifying fluorescence recovery after photobleaching (FRAP) using microtubule probes.

Microtubule turnover requires assembly of new fluorescent tubulins into the bleached area. The rate of recovery of the microtubule signal after photobleaching was similar in isogenic control and mutant neurons (**Figure 4H-I**). Thus, it is unlikely that lysosomal trafficking defects are driven by microtubule alterations in *MAPT* p.R406W neurons.

The positioning and trafficking of lysosomes rely on a complex interplay of interactions between microtubule motors and motor-associated proteins, such as JIP3, ARL8, RILP ^44, 45^. To determine whether *MAPT* p.R406W impacts motor adaptor proteins, we evaluated expression of motor adaptor genes in our transcriptomic data (**Figure 4J**). Among the candidate motor-associated genes, only *JIP3* was significantly upregulated in *MAPT* p.R406W neurons compared with isogenic controls (**Figure 4J**). JIP3 protein was also significantly elevated in mutant neurons (**Figure 4K-L**). The motility of the most mature population of autophagic vesicles is regulated by the motor-interacting protein JIP3 ^22, 44^. As autophagosomes and lysosomes travel along axons, they acquire multiple motor-interacting proteins that sequentially regulate dynein activity and lysosome trafficking, depending on their maturation status and location within axons (**Figure 4M**) ^22^. Interestingly, lysosome distribution in neurites is sensitive to JIP3 levels: JIP3 knockout and JIP3 overexpression result in axonal swellings ^46^, suggesting that JIP3-mediated functions are tightly regulated by JIP3 expression.

### *MAPT* p.R406W neurons exhibit impairment in autophagy

Proper lysosome positioning is essential for autophagy ^47^. This process starts with the engulfment of organelles into an autophagosome, which subsequently fuses with a lysosome (**Figure 5A**). This process is pivotal to stymie neurodegenerative disease, as it facilitates the recycling of dysfunctional proteins ^34^. Transcriptomic analyses pointed to the dysregulation of genes enriched in phagosome maturation, the autophagosome, and regulation of autophagy (**Figure 3A**). To determine whether *MAPT* p.R406W contributes to defective autophagic flux, we assessed the overall abundance of autophagosomes in *MAPT* p.R406W and isogenic controls using CYTO-ID (**Figure 5B**). Mutant neurons produced significantly more autophagosomes compared to isogenic controls (**Figure 5B-C**). To determine whether this change was due to an increase in autophagosome biogenesis or reduced autophagic flux, neurons were treated with rapamycin (to promote autophagosome biogenesis) and chloroquine (to block autophagic flux; **Figure 5B**). Compared to DMSO-treated neurons, rapamycin and chloroquine treatment significantly enhanced CYTO-ID intensity and CYTO-ID positive vesicle density in both mutant and isogenic control neurons (**Figure 5B-D**). In rapamycin and chloroquine treated neurons, *MAPT* p.R406W and control neurons exhibited similar CYTO-ID intensity and density of CYTO-ID positive vesicle (**Figure 5B-D**). LC3B sits on the autophagosome membrane where it serves as a marker of autophagosome initiation and maturation ^48^. Consistent with our CYTO-ID findings, LC3B was significantly elevated in mutant neurons (**Figure 5E, 5G**). Together, these observations suggest that *MAPT* p.R406W increases autophagosome accumulation and maturation at baseline.

**Figure 5:**
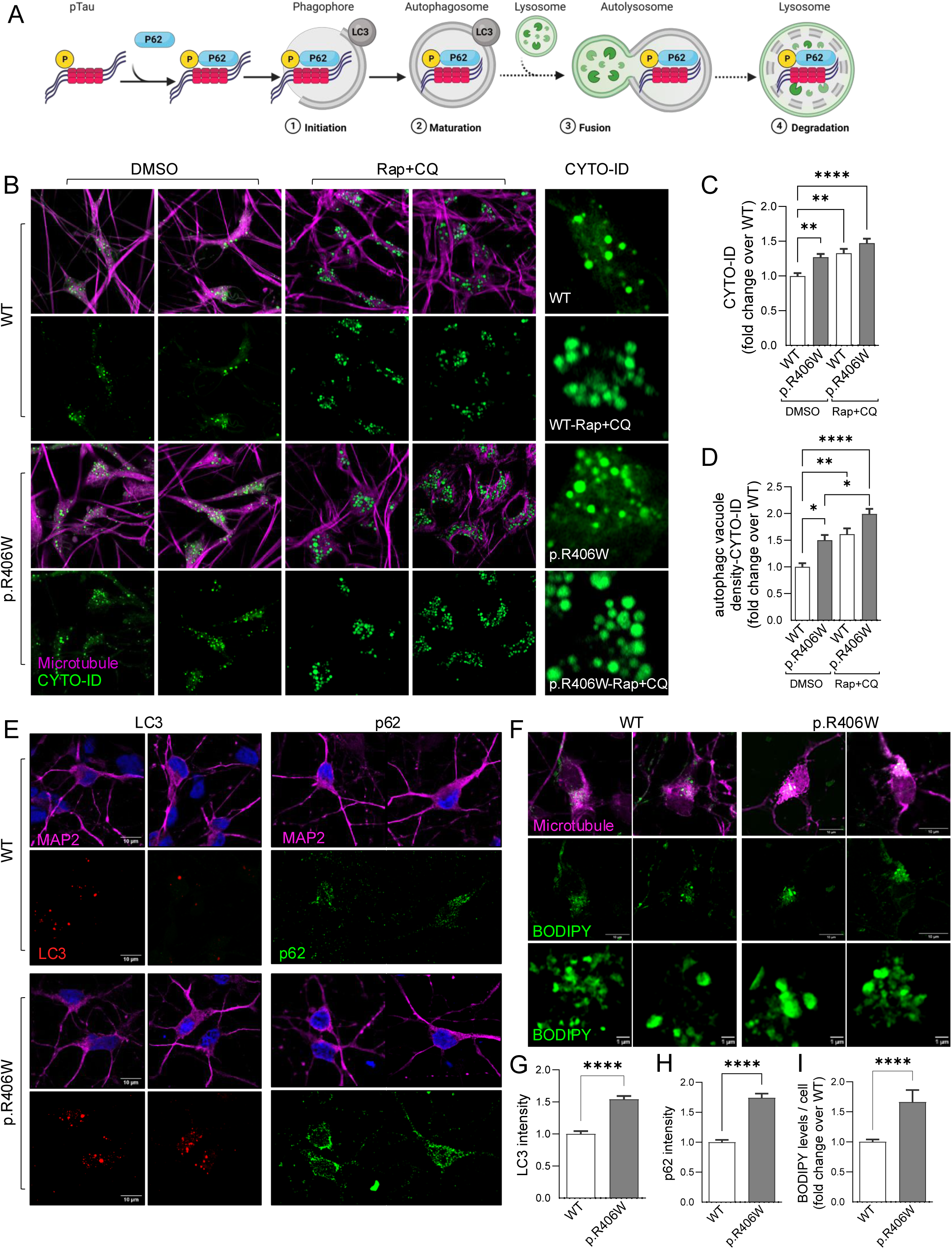
*MAPT* p.R406W neurons exhibit impairment in autophagy. *MAPT* p.R406W and isogenic control neurons probed for autophagic markers at DIV14. A. Diagram of major steps of macroautophagy and autophagic fusion with lysosomal structures. B. Autophagosomes were monitored in live neurons using CYTO-ID (green) along with MAP2 to capture microtubule (magenta). Scale bar 10µm. Rapamycin (Rap, 500nM) and Chloroquine (CQ, 20uM) treatments were used as positive control (16hrs). Right panel, magnified CYTO-ID positive vesicles (green). Scale bar 1µm. C. Quantification of CYTO-ID mean fluorescence intensity per cell. Data are represented as mean ± SEM (total number of cells quantified for WT n=26, WT-Rap+CQ n=29, p.R406W n=34, pR406W-Rap+CQ n=29 from 3 independent experiments). Kruskal-Wallis test followed by Dunn’s test, **, p<0.005,****, p<0.0001. D. Quantification of CYTO-ID mean density (number of autophagic vacuoles, CYTO-ID+ puncta per cell). Data are represented as mean ± SEM (total number of cells quantified for WT n=17, WT-Rap+CQ n=20, p.R406W n=17, pR406W-Rap+CQ n=19 from 3 independent experiments). Kruskal-Wallis test followed by Dunn’s test. * *p<0.05, ** p<0.001, **** p<0.0001.* E. Neurons were stained for MAP2 (magenta), LC3B (red) and p62 (green) in p.R406W neurons and isogenic controls, Scale bar 10µm.F. Neurons immunostained with MAP2 (magenta) and BODIPY (lipid droplets, green). Scale bar 10µm. Lower panels, magnified BODIPY-positive neutral lipids stained in green. Scale bar 1µm. G. Quantification of LC3B mean intensity per cell (number of cells quantified for *MAPT* WT n=36 and p.R406W n=47 ). Mann-Whitney *U* test. ****, *p*<0.0001. H. Quantification of p62 mean intensity per cell (number of cells quantified for *MAPT* WT n=33, and p.R406W n=36) Mann-Whitney *U* test. ****, *p*<0.0001. I. Quantification of relative BODIPY mean intensity per cell (number of cells quantified for *MAPT* WT n=51 and p.R406W n=61). Mann-Whitney *U* test. ****, *p*<0.0001. G-I. Data are mean ± SEM and are quantified from 3 independent experiments.

Autophagosomes undergo fusion with lysosomes, where the vesicle cargo goes on to be degraded by lysosomal hydrolases ^49^. P62 (also known as sequestosome-1 or SQSTM1) binds to cargo, facilitating cargo degradation within autophagosomes ^50, 51, 52^. When autophagic degradation is disrupted or inefficient, p62 levels increase ^52^. *MAPT* p.R406W neurons exhibited significantly more p62 than isogenic controls (**Figure 5E** and **5H**). Thus, despite an increase in autophagosomes (CYTO-ID and LC3B), cargo was not degraded efficiently in mutant neurons.

To determine whether the observed defects in autophagy impact cargo degradation more broadly, we evaluated neutral lipids. While these lipid droplets are not harmful by themselves, they can represent disrupted lipid processing and distribution ^53^. To determine whether *MAPT* p.R406W impacts lipid droplet biogenesis, we used BODIPY 493/503. We observed significantly more lipid droplet content in mutant neurons relative to controls based on the overall BODIPY fluorescence per cell (**Figure 5F, 5I**). These data suggest that *MAPT* p.R406W induce defects in autophagy that have a broad impact on cargo, which ultimately promotes lipid dysregulation.

### Autophagy enhancer G2-567 rescues Tau defects in *MAPT* p.R406W neurons

Given our findings of Tau accumulation and autophagy defects in *MAPT* p.R406W neurons, we sought to determine whether pharmacological enhancement of autophagy could reverse Tau accumulation. G2-567 is a small molecule that enhances autophagy. The 567 analog is more potent and stable than the 115 analog, which has been shown to promote autophagic degradation of α1-antitrypsin variant Z and huntingtin in mammalian cells ^54, 55^.

*MAPT* p.R406W neurons and isogenic controls were treated with G2-567 (0.5 µM for 14 days) or DMSO vehicle control. G2-567 significantly reduced pTau levels in mutant neurons compared to controls (**Figure 6A-B**). To determine whether G2-567 impacts Tau accumulation in LAMP1+ vesicles, we imaged treated neurons using Airyscan super resolution microscopy. In G2-567-treated mutant neurons, the percentage of lysosomes free of Tau was significantly higher than control treated mutant neurons (p.R406W-DMSO: 32.8% vs. p.R406W-G2-567: 67.1%; **Figure 6C-D**). Enlargement of LAMP1+ vesicle size in *MAPT* p.R406W neurons was restored to sizes observed in controls upon G2-567 treatment (**Figure 6E-F**). In fact, neurons treated with G2-567 displayed a significant reduction in lysosomal size in both mutant and isogenic control neurons (**Figure 6E-F**). Our findings demonstrate that treating mutant neurons with an autophagy enhancer, G2-567, rescues pTau levels, increases Tau-free lysosomes, and reduces lysosome size (**Supplemental Figure 6**).

**Figure 6:**
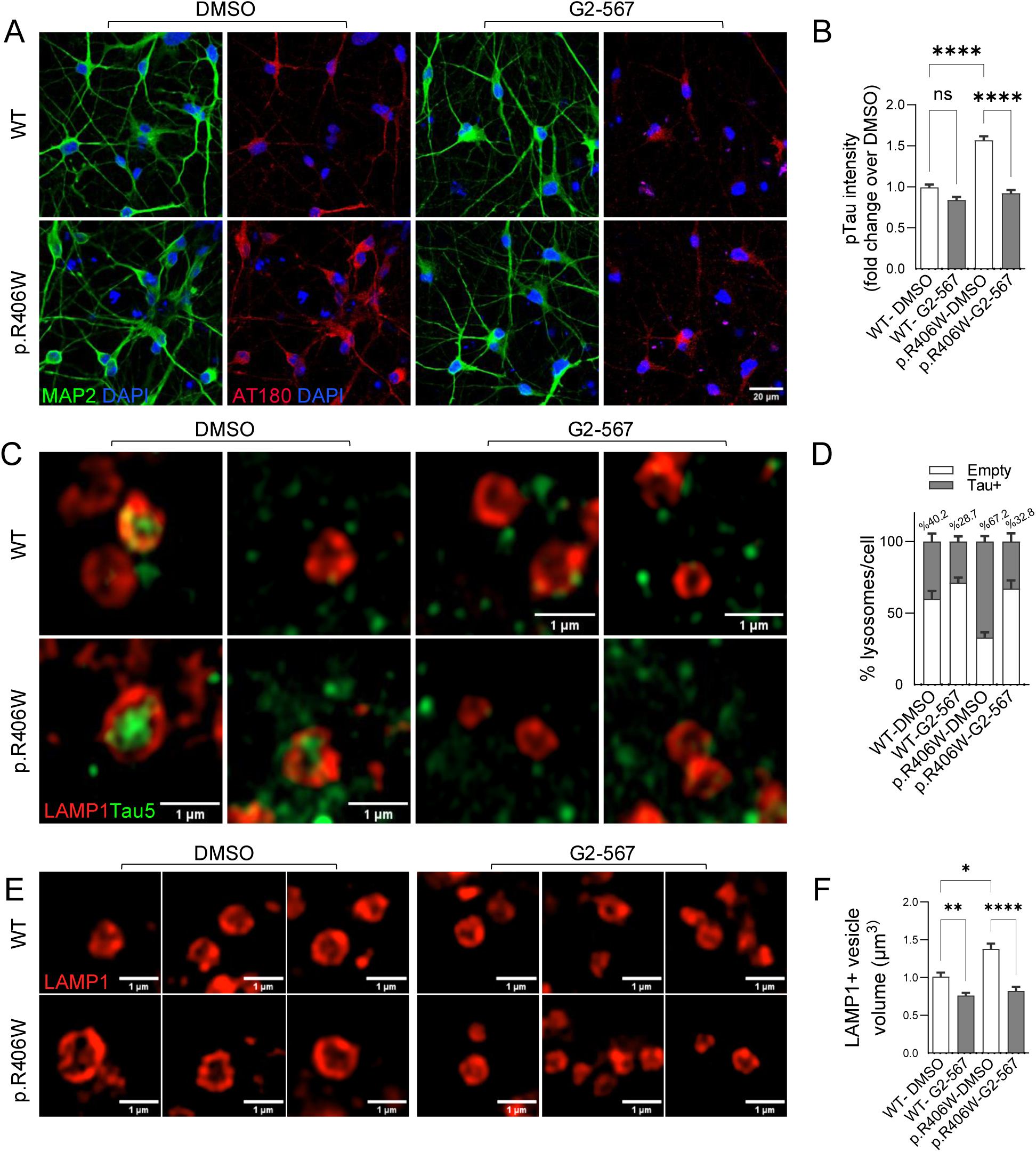
Autophagy enhancer G2-567 rescues tau defects in *MAPT* p.R406W neurons. *MAPT* p.R406W and isogenic control neurons were treated with G2-567 (0.5µM) and DMSO for 14 days beginning on DIV7, fixed on D21. A. Neurons (MAP2, green) stained for pTau (AT180, red). Scale bar 20µm. B. Quantification of pTau mean intensity per cell (WT n=69, WT-G2-567 n=69, p.R406W n=59, pR406W-G2-5567 n=56). Data are represented as mean ± SEM. Kruskal-Wallis test followed by Dunn’s test. ****, *p*<0.0001. C. LAMP1 vesicles (red) co-stained with total Tau (Tau5, green), in *MAPT* p.R406W and isogenic control neurons after treatment with G2-567. Scale bar 1µm. D. Quantification of the percentage of lysosomes free of total Tau (empty, white) or co-localized with total Tau (Tau positive, gray). Data are represented as mean ± SEM. ANOVA mixed model multiple comparison. p.R406W-DMSO: Empty vs. p.R406W-G2-567: Empty *, p= 0.0171. E. LAMP1+ vesicles (red) in *MAPT* p.R406W and isogenic control neurons after treatment with G2-567. Scale bar 1µm. F. Quantification of lysosome volume. Kruskal-Wallis test followed by Dunn’s test, *, *p*=0.0109, \*\**p*=0.0036, ****, *p*<0.0001. Data are representative of three independent experiments.

### Autophagy enhancer G2-567 increases autophagic flux in *MAPT* p.R406W neurons without rescuing motility defects

Next, we sought to determine whether treatment with G2-567 rescues the motility and autophagic defects in *MAPT* p.R406W neurons. G2-567 treatment did not significantly alter the number of LAMP1+ vesicles compared to vehicle treatment in mutant or control neurons (**Figure 7A-B**). G2-567-treated mutant neurons also maintained a significant increase in distally positioned lysosomes compared to vehicle treated controls (**Figure 7A** and **7C**). To determine whether G2-567 restores the expression of the dysregulated motor adaptor protein JIP3, we evaluated JIP3 immunofluorescence signal intensity in *MAPT* p.R406W and isogenic control neurons. JIP3 levels remained significantly elevated in G2-567 treated mutant neurons compared with vehicle controls (**Figure 7D-E**). Thus, G2-567 treatment does not restore lysosomal positioning, lysosomal number, or motor adaptors in mutant neurons.

**Figure 7:**
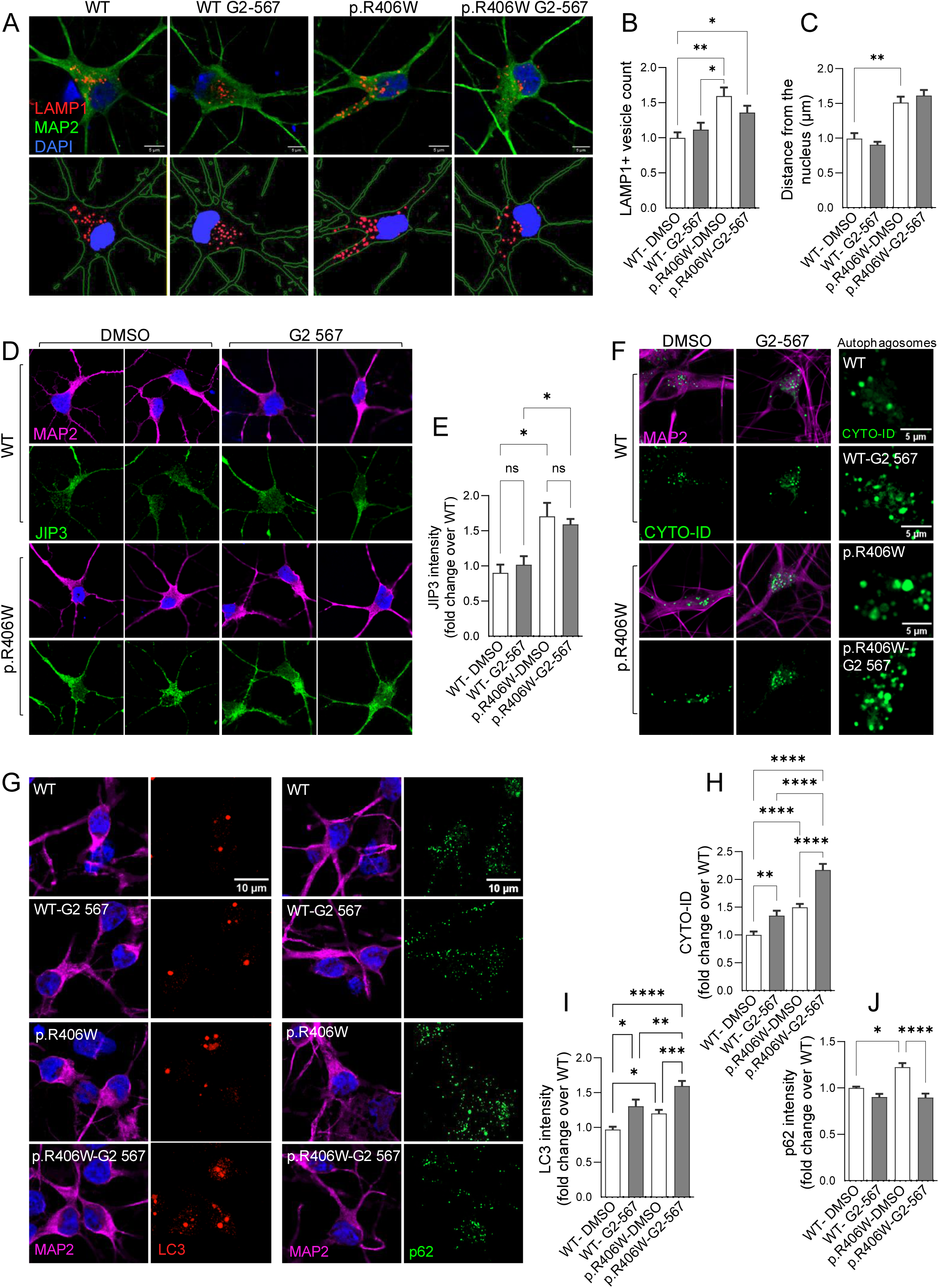
Autophagy enhancer G2-567 increases autophagic flux in *MAPT* p.R406W neurons without rescuing motility defects. *MAPT* p.R406W and isogenic control neurons were treated with G2-567(0.5µM) or DMSO for 14 days beginning on DIV7 and fixed on DIV21. Neurons were then immunostained for lysosomal positioning and autophagy markers. A. Neurons were immunostained for MAP2 (green) and LAMP1 (red). Lower panel, 3D reconstruction by Imaris. Scale bar 5µm. B. Quantification of lysosome density (WT n=16, WT-G2-567 n=14, p.R406W n=17, p.R406W-G2-567 n=19). Kruskal-Wallis test followed by Dunn’s test. *, *p<0.05*, *\*\*, p<0.0017.* C. Quantification of lysosome distance from the nucleus (measurement of shortest distance of LAMP1+ vesicles from DAPI, quantified by immaris 3D rendering. Number of vesicles quantified for WT n=181, WT-G2-567 n=219, p.R406W n=319, p.R406W-G2-567 n=318). Kruskal-Wallis test followed by Dunn’s test, WT-DMSO vs. p.R406W-DMSO, **, p= 0.0011, p.R406W-DMSO vs. p.R406W-G2-567, p= 0.3235. D. Neurons were immunostained with MAP2+ (magenta) and JIP3 (green). E. Quantification of JIP3 total mean intensity (n=1 experiment total 22 cells). ANOVA, *, *p<0.05,* p.R406W-DMSO vs. p.R406W-G2-567, *p=*0.9908. F. *MAPT* p.R406W and isogenic control neurons from DMSO control and G2-567-treated cells were probed for autophagosomes using CYTO-ID (green) and MAP2 (magenta). Right panel, magnified CYTO-ID positive vesicles (green). Scale bar 5µm. G. Neurons treated with G2-567 were immunostained for MAP2 (magenta), LC3B (red) and p62 (green). Scale bar 10µm. H. Quantification of CYTO-ID mean density. Data is representative of 3 independent experiments. (Total number of cells quantified for WT n=71, WT-G2-567 n=79, p.R406W n=79, p.R406W-G2-567 n=78). Kruskal-Wallis test followed by Dunn’s test, **, *p<0.005, ****, p<0.0001.* Data from 3 independent experiments. I. Quantification of LC3B mean intensity per cell. (Total number of cells quantified for WT n=70, WT-G2-567 n=43, p.R406W n=68, p.R406W-G2-567 n=68). Kruskal-Wallis test followed by Dunn’s test, * *p*<0.05, **, *p*<0.01, ****, *p*<0.0001. J. Quantification of p62 mean intensity. (Total number of cells quantified for WT n=26, WT-G2-567 n=32, p.R406W n=32, p.R406W-G2-567 n=26). Kruskal-Walli’s test. *, *p*<0.05, ****, *p*<0.0001. Data in I and J are from 4 independent experiments. All data are mean ± SEM.

To understand the impact of G2-567 on autophagy, we measured autophagic flux in mutant and control neurons after G2-567 treatment. We observed a significant increase in the CYTO-ID signal, representing more autophagic vacuoles, with G2-567 treatment in both mutant and control neurons (**Figure 7F** and **7H**). This finding is consistent with the proposed mechanism of action of G2-567. Autophagosome surface marker, LC3B, was also significantly elevated in G2-567 treatment compared with vehicle controls in both mutant and isogenic control neurons (**Figure 7G** and **7I**). Thus, G2-567 enhances autophagic flux in mutant and control neurons. To determine whether the enhanced autophagic flux leads to productive cargo degradation, we measured p62 levels. Treatment of *MAPT* p.R406W neurons with G2-567 resulted in a significant reduction of p62 (**Figure 7G** and **7J**). G2-567 treatment did not alter p62 levels in isogenic control neurons (**Figure 7G** and **7J**). Thus, G2-567 enhanced productive autophagy in mutant neurons. We replicated the same phenotype in a second donor line, further confirming the robustness of our findings (**Supplemental Figure 7**). Together, our findings suggest that G2-567 rescues pTau levels and Tau accumulation in lysosomes by enhancing autophagy without rescuing motility defects.

## Discussion

Here, we provide the first high-dimensional visualization of endogenous Tau degradation in human neurons. We discovered key differences in the way human neurons handle Tau and pTau in the lysosome. These differences in Tau and pTau are exaggerated in neurons expressing *MAPT* p.R406W, where there is an overall degradation deficiency. We find that *MAPT* p.R406W neurons exhibit broad transcriptomic and defects autophagy and lysosomal, which contribute to the defects in degradative capacity. Beyond Tau and pTau, the *MAPT* p.R406W mutation is sufficient to disrupt p62 and neutral lipid degradation. We were able to rescue mutant tau-induced defects in Tau and pTau degradation by restoring productive autophagy without rescuing lysosomal trafficking defects using an autophagy enhancing drug, G2 series. Together, our findings create a more complete picture of the direct impact of tau mutations to the degradative capacity of human neurons and provide evidence that we can de-couple lysosomal trafficking and degradative capacity to therapeutically target protein clearance.

Our study suggests that Tau degradation is driven by an interplay between neuronal trafficking defects and faulty autophagy pathways. In lysosomal storage disorders, cells respond to inefficient degradative machinery by upregulating macroautophagy components ^56^. We reported enhanced autophagic flux coupled with delayed cargo degradation in *MAPT* p.R406W neurons, which is reminiscent of lysosomal storage disorders. Importantly, we discovered that the chemical enhancement of autophagy rescued pTau levels and lysosomal phenotypes without restoring the trafficking defects – effectively decoupling motility from degradation.

Post-translational modifications on pathogenic proteins, such as Tau, can directly and indirectly impair autophagy pathways ^13, 57, 58^. We discovered that the presence of mutant Tau (p.R406W) was sufficient to induce several changes in the association of Tau with the lysosome. Neurons expressing mutant Tau had fewer lysosomes lacking Tau or pTau protein, suggesting that degradation of Tau is stalled. These defects were restored upon correction of the mutant allele by CRISPR/Cas9. Additionally, we discovered that phosphorylated Tau accumulates at the lysosomal membrane in mutant neurons. While phosphorylation of Tau is required for normal function, hyperphosphorylation is a key feature of Tau aggregates. Thus, as has been suggested for acetylation of Tau, pTau may directly impair autophagic machinery early in a manner that ultimately promotes Tau aggregation ^27^.

Precise regulation of lysosome localization and motility are essential for maximal degradative capacity in neurons ^18, 20, 23^. Neurons show spatially restricted autophagosome biogenesis, which is mostly enriched at distal ends of axons in both growing and synaptically connected neurons ^59^. Distal lysosomes are largely devoid of degradative enzymes and need to undergo retrograde transport to the neuronal soma for effective cargo degradation ^42, 60, 61^. Late endosomes predominantly undergo long-range retrograde transport while simultaneously acidifying and maturing into lysosomes, and ultimately fusing with mature lysosomes enriched in the cell body ^62^. For efficient degradation, endolysosomal vesicles must reach the soma where the vesicles fuse with lysosomes ^17, 63^. JIP3 silencing has been shown to reduce lysosome association with dynein motors, leading to failure in retrograde trafficking. Loss-of-function mutants in JIP3 display lysosome accumulations in axonal terminal swellings, and an increase in pTau protein ^46^. We report that *MAPT* p.R406W causes impairment in retrograde transport that is mediated by the motor molecule JIP3. Thus, the observed accumulation of pTau in *MAPT* p.R406W neurons may be driven by defects in lysosomal trafficking due to faulty motor associated proteins, which presents a potential therapeutic target. However, rescuing JIP3 levels is challenging - prior studies illustrate that JIP3 overexpression also induces lysosomal and axonal defects ^64^.

Alterations in intracellular lipid content, such as in metabolic disorders, can significantly affect the fusion step of macroautophagy and influence the overall activity of this intracellular proteolytic pathway ^65^. Membrane lipids are broken down into fatty acids that can serve as a source of energy production or be packaged into lipid droplets, acting as reservoirs for stored lipids destined for future utilization. It is common for patients with neurodegenerative disorders, including FTD, to have abnormal levels of lipid droplets in their nerve cells ^66^. It has also been demonstrated how upregulated lipid synthesis and downregulated lipid turnover lead to pathological lipid accumulation in both neurons and glia in multiple tauopathy model systems. Here we showed that mutant neurons have a greater lipid droplet content compared to the WT control neurons, indicating a more general neuronal metabolic dysregulation.

Derivatives of glibenclamide, including G2-115, have been reported to increase autophagy in an mTOR independent manner without altering insulin secretion ^67^. G2-115 treatment have been shown to increase LC3, reduce p62, and reduce levels of α1-antitrypsin Z (ATZ) and Huntingtin proteins in *C. elegans* models, directly converted human neurons and animal models ^55, 67^. G2-115 acts as an autophagy enhancer by reducing the interaction between RCAN1 and calcineurin. This stimulates calcineurin activity and promotes the translocation of transcription factor EB (TFEB) into the nucleus where it regulates expression of genes associated with lysosomal biogenesis ^54, 55^. Calcineurin is a phosphatase that acts on Tau ^68^. Inhibition of calcineurin activity promotes Tau phosphorylation and secretion from the cell ^69^. Similarly, we found that enhancing autophagy with an analog of G2, G2-567, was sufficient to reduce pTau levels and to increase the proportion of lysosomes free of Tau in mutant neurons. Thus, G2-567 and its derivates promote broad protein degradation *in vitro* and *in vivo*.

Overall, our study unveils an interplay between mutant Tau and its impact on the cellular clean up system, suggesting that *MAPT* p.R406W may interfere with the normal physiological clearance of Tau. Together, our findings suggest that *MAPT* p.R406W may be sufficient to cause impaired lysosomal transport and autophagy, and that restoring autophagy is sufficient to rescue pTau levels and degradative capacity of the mutant neurons and in a stem cell model of tauopathy. More than 50 mutations occur in the *MAPT* gene and cause FTLD-Tau ^70^. So, the extent to which these phenotypes are shared across different tau mutations must be investigated in the future.

## Methods

### iPSC lines

iPSCs from a *MAPT* p.R406W carrier (F11362.1 or FA14530.1) and the CRISPR/Cas9 edited isogenic control (F11362.1Δ1B06 or FA14530.1Δ3G12) were previously described ^71^. To facilitate the generation of excitatory neuronal cultures, a Tet-ON 3G-controlled NGN2 transgene was integrated into the AAVS1 locus of the iPSC lines described above using TALENs ^30, 72^: *MAPT* p.R406W donor line (iPSC F11362.1.CRISPRiΔ5A01 .NGN2Δ1B08 and iPSC FA14-530.1.NGN2Δ1C05) and isogenic control iPSC F11362.1Δ1C11.CRISPRiΔ2D10.NGN2Δ1C07 and iPSC FA14-530.1Δ3G12.NGN2Δ1E04). Engineered cell lines were verified using Sanger sequencing for the *MAPT* mutation and the NGN2 cassette, karyotyping for chromosomal stability, and immunocytochemistry for pluripotency markers (**Supplemental Figure 1-2**). Cell lines were maintained in mTesR medium (STEMCell Technologies, 85850) on Cultrex PathClear Basement Membrane Extract, Type II (R&D Systems, 3532-010-02). The cell lines were confirmed to be free of mycoplasma.

### iPSC differentiation

*MAPT* p.R406W iPSC and WT, isogenic controls with the integrated NGN2 transgene were differentiated into neural progenitor cells (NPCs) as previously described ^71^. Briefly, iPSCs were plated at density of 65,000 cells per well in Neural Induction Media (STEMdiff SMADi Neural Induction Kit STEMCell Technologies, 08581) in a 96-well v-bottom plate to form neural aggregates. After 5 days, neural aggregates were passaged onto Poly-L-Ornithine (PLO, R&D Systems, 3436-100-01) and laminin-coated (R&D Systems, 3400-010-03) plates to form neural rosettes. Neural rosettes were isolated after 5 days in culture by selection using STEMdiff Neural Rosette Selection Reagent (STEMCell Technologies, 05832). The resulting NPCs were cultured on PLO and laminin-coated plates, and terminal differentiation was initiated with the addition of cortical maturation medium (BrainPhys Neuronal Medium, STEMCell Technologies, 05790) supplemented with B27 (Gibco, 17504001), BDNF (Peprotech, 450-02-100UG) (10 ng/mL), GDNF (Peprotech, 450-10-100UG) (10 ng/mL), cAMP (Sigma, D0627-1G), GlutaMax (Thermo Scientific, 35050061) N2 supplement (Gibco, 17502001) and doxycycline (Sigma, D5207-10G) (2 mg/mL). Neurons typically show neuronal identity 7 days after plating based on immunocytochemistry for β-tubulin III (Tuj1) and microtubule-associated protein 2 (MAP2), a marker of mature neuronal dendrite (**Supplemental Figure 3**). Neurons were analyzed 14 or 21 days after doxycycline induction.

### Antibodies

Primary antibodies used for immunostaining included: anti-MAP2 (Abcam, ab5392), anti-LAMP1 (D2D11; Cell signaling, 9091), anti-Tau5 (total Tau; generously provided by Dr. Lester Binder), anti-AT180 (pTau-Thr23; Thermo Fisher Scientific, MN1040), anti-AT8 (pTau-Ser202, Thr 205; Thermo Scientific, MN1020B), anti-JIP3 (Thermo Fisher, PA5-59728), anti-LC3B (Cell Signaling, 2775), anti-SQSTM1 / p62 (Cell signaling, D5L7G 88588S), anti-Synapsin I (Millipore, AB1543), Cathepsin D (C-5; SantaCruz, sc-377124), anti-Beta III Tubulin (Tuj1; Promega, G712A), and DAPI (Sigma, D-9542 ). The secondary antibodies included: Alexa-488 Goat anti-mouse IgG (H+L; Thermos Scientific, A-11001), Alexa-488 Goat anti-rabbit IgG (H+L; Thermos Scientific, A-11008), Alexa-568 Goat anti-mouse IgG (H+L; Thermos Scientific, A-11004), Alexa-568 Goat anti-rabbit IgG (H+L; Thermos Scientific, A-11011), Alexa-647 Goat anti-mouse IgG (H+L; Thermos Scientific A-32728), Alexa-647 Goat anti-rabbit IgG (H+L; Thermos Scientific A-32733), Alexa Fluor™ Plus 647 Goat anti-Chicken IgY (Thermos Scientific A-32933), and Alexa-405 Goat anti-chicken IgY (Abcam, ab175674) (**Supplemental Table 1**).

### Immunostaining

iPSC-derived neurons plated on coverslips were washed and fixed with 4% paraformaldehyde (Electron Microscopy Sciences, 15710) for 20 minutes at room temperature (RT), and permeabilized with 0.3% Triton X-100 in PBS for 5 minutes at room temperature (RT). Cells were blocked with 2% FBS,1% BSA, and 0.1% saponin in PBS and incubated with primary antibodies at 4°C overnight. After washing in PBS, neurons were incubated with secondary antibodies for 1 hour at RT. Cells were incubated with DAPI (Sigma, D-9542) for 5 minutes then washed with PBS. Coverslips were then mounted using Fluoromount-G (Southern Biotechnology, 00-4958-02). Confocal microscopy was performed with LSM 980 confocal microscope (AiryScan, Zeiss). Each image reflects a Z-stack of 5–12 images each taken at 0.2 μm depth intervals. Samples were imaged using identical acquisition parameters for direct comparison.

### Monitoring Acidic Vesicle Motility in Live Cells

To evaluate motility of acidic vesicles, iPSC-derived neurons were cultured in live-imaging dishes pre-coated with poly-L-ornithine and laminin (35mm glass-bottom dish #1.5H, Cellvis, D35-20-1.5H) at a density of 100,000-150,000 cells/dish. Neurons were probed for microtubules by ViaFluor® dye (Viafluor^®^ 647, Biotium, 70063, 1x in media, 30minute incubation) to define cell boundaries. Cells were then incubated with 50nM LysoTracker Red D99 (Invitrogen, L7528), for 10 minutes. Super-resolution confocal microscopy (Zeiss LSM 980 Multiphoton Airyscan 2; Carl Zeiss Ltd, Cambridge, UK) was used with a 40x or 63x objective lenses to image neurons or record and track LysoTracker-positive vesicles in time-lapse mode for up to 5 minutes. Each assay was repeated at least 3 times from independent cultures. Only modest bleaching was observed during the experiment. Cells showing signs of damage (pearling or blebbing) were discarded. Lysosomal motility/trafficking was measured in a semi-automatic manner using Imaris (Bitplane) by tracking LysoTracker-positive puncta over time, using the 3D time-lapse tracing module. The lysosomal tracks in neurites were visualized in order to quantify acidic vesicle velocity and vesicle displacement length. Acidic vesicles within the field were detected using the spot function, and tracks were semi-automatically created using the Brownian motion algorithm in Imaris.

### Monitoring Autophagy in Live Cells

For autophagy assays, iPSC-derived neurons were cultured in live-imaging dishes pre-coated with poly-L-ornithine and laminin. Neurons were incubated with CYTO-ID (1:1000; Enzo 175-0050) for 20 minutes based on manufacturer’s recommendations, and live imaging was performed at 37 °C in a humidified and CO_2_-controlled chamber. Neurons were also probed for microtubule by ViaFluor® dye (Viafluor^®^647, Biotium, 70063) to define cell boundaries. Images were acquired by super-resolution confocal microscope (Zeiss LSM 980 Airyscan). CYTO-ID selectively probes the autophagosome membrane and vesicles appear as round-shaped structures detectable under fluorescence excitation/emission of 495/519 nm. Density and intensity of CYTO-ID-positive autophagic vacuoles were quantified within the neurons defined by microtubule signal using Fiji ImageJ.

### Monitoring Lipids in Live Cells

Levels of intracellular neutral lipids were quantified by adding BODIPY^TM^ FL C_16_ (Thermo Scientific D3821, final concentration 10 μg/ml in PBS) for 20 minutes at RT to iPSC-derived neurons. Live imaging was then immediately performed at 37 °C in a humidified and CO_2_-controlled chamber. Neurons were also probed for microtubule by ViaFluor® dye (Viafluor^®^ 647, Biotium, 70063) to define cell boundaries. Images were acquired using a super-resolution confocal microscope (Zeiss LSM 980 Airyscan). BODIPY^TM^ FL C_16_ probes neural lipid storage compartments detectable under fluorescence excitation/emission of 505/512 nm. BODIPY intensity was measured within the microtubule signal using Fiji ImageJ.

### Image Quantification of Tau/pTau Localization in Lysosomes

Tau (Tau5) and pTau (AT8) localization was quantified in LAMP1-positive vesicle using Fiji ImageJ. The signal intensity (Gray Value) of Tau or pTau within each LAMP1-positive vesicle was measured. A single line spanning the LAMP1-positive vesicle was used to define the plane for quantification of Tau or pTau signal intensity. Intensity curves were then generated from the selected pixels. Total Tau and pTau intensity measurements were performed using Fiji as total mean fluorescence intensity per cell using structural signals from MAP2 as the cell boundary. Measurements were corrected for background fluorescence resulting in the Corrected Total Cell Fluorescence (CTCF) values.

### Image Quantification of Lysosome Characterization

Lysosome characterization analysis was performed using LAMP1 antibody (D2D11; Cell signaling, 9091), on 60x confocal images acquired in Z-stacks. Lysosomes were 3D reconstructed in Imaris for measurements of lysosome count, volume, and distance from soma by first detecting the LAMP1+ puncta using the Imaris spot detection module. We then measured the spot density and volume as well as the distance of the detected spot (lysosome) to the nucleus, as a measurement for lysosomal distribution.

### Image Quantification of Neuronal Morphology

Neuronal morphometric analysis was performed on 40x confocal images acquired in Z-stacks. Images were 3D reconstructed in Imaris software using filament module and were semi-automatically analyzed for dendritic arborization and length as well as Sholl analysis(Imaris Sholl Analysis XTension, Sholl sphere resolution 1um).

### Fluorescence Recovery After Photobleaching (FRAP)

FRAP experiments were performed using the FRAP module on the confocal microscope (Zen software). Neurons (DIV14) were incubated with the microtubule probe Viafluor® 488 (Biotium, final concentration: 1uM) for 15 minutes at RT. Neurons were then imaged for 20 seconds (pre-bleach phase). A circular region of interest (ROI) was selected for photobleaching along the neurites. Photobleaching was performed at 50% laser power and 50 iterations. Images were acquired at 2 seconds intervals over a period of 5 minutes following the recovery of the fluorescence intensity in the selected ROI. FRAP recovery curves were generated using Fiji with normalization to the initial pre-bleach value of fluorescence intensity.

### G2 Treatment

The small molecule G2 was identified by a phenotypic screen using a *C.elegans* model which expresses the misfolded α1-antitrypsin Z variant (ATZ) ^73^ and later shown to mediate autophagic degradation of ATZ in mammalian cell line models ^67^. This analog has been subjected to extensive chemical modification to derive the G2-567 compound. *MAPT* p.R406W and isogenic control iPSC-neurons were treated with 0.5 µM G2-567 (resuspended in DMSO) on DIV7, a time point where neurons are positive for neuronal markers MAP2 and Tuj1 (**Supplemental Figure 3**). Neurons were treated every 2 days for 14 days. Neurons were then fixed on DIV21 for immunofluorescent staining or live imaging as described above.

### RNAseq

To evaluate the impact of *MAPT* p.R406W on molecular pathways, we analyzed RNAseq data from *MAPT* p.R406W and isogenic control iPSC-derived neurons that were previously described ^33^. Briefly, DNA libraries of individual samples were constructed using the TruSeq Stranded Total RNA Sample Prep with Ribo-Zero Gold kit (Illumina) and then sequenced by Illumina HiSeq 4000 Systems Technology with a read length of 1×150 bp, and an average library size (mapped reads) of 43.2 ± 9.5 million reads per sample. The average percentage of mapped reads to GRCh38 was 94.0% ± 2.0. STAR (v.2.6.0) ^74, 75^ was used to align the newly generated RNA sequences to the human reference genome GRCh38.p13 (hg38). Salmon (v. 0.11.3) ^76^ was used to quantify the expression of the genes annotated within the human reference genome used in this project (GRCh38.p13). Protein coding genes were selected for further analyses. Differential gene expression analysis was performed between samples carrying the mutation and controls was calculated using the DESeq2 (v.1.22.2) package ^77^.

## Acknowledgments

We thank the research subjects and their families who generously participated in this study. We thank Dr. Binder who generously provided the Tau5 antibodies. We thank David Kast and Torri Ball for thoughtful discussions. We thank the Genome Engineering & Stem Cell Center (GESC@MGI) at Washington University in St. Louis School of Medicine for iPSC reprogramming services. This work was supported by access to equipment made possible by the Hope Center for Neurological Disorders, the Neurogenomics and Informatics Center, and the Departments of Neurology and Psychiatry at Washington University School of Medicine. Confocal images were generated on a Zeiss LSM 980 Airyscan Confocal Microscope which was purchased with support from NIH grant number S10 MH126964 through the use of Washington University Center for Cellular Imaging (WUCCI)-Neuro supported by Washington University School of Medicine, the Children’s Discovery Institute of Washington University and St. Louis Children’s Hospital (CDI-CORE-2015-505 and CDI-CORE-2019-813), and the Foundation for Barnes-Jewish Hospital (3770 and 4642). Funding provided by the National Institutes of Health (P30 AG066444, R01 AG056293, RF1 NS110890, U54 NS123985), Hope Center for Neurological Disorders (CMK), Rainwater Charitable Organization (CMK), Farrell Family Fund for Alzheimer’s Disease (CMK), and UL1TR002345. Diagrams were generated using BioRender.com.

## Supplemental Figure Legends

**Supplemental Table 1:** Antibodies used for immunofluorescence staining.

**Supplemental Figure 1: iPSC characterization for donor pair 1.** iPSC characterization of *MAPT* p.R406W donor line (iPSC F11362.1.CRISPRiΔ5A01.NGN2Δ1B08) and isogenic control iPSC F11362.1Δ1C11.CRISPRiΔ2D10.NGN2Δ1C07). Each iPSC line used in the study was verified by Sanger sequencing. Karyotyping was performed to confirm chromosomal stability. Immunocytochemistry was performed to confirm expression of pluripotency markers as well as NGN2 plasmid sequencing for integration confirmation.

**Supplemental Figure 2: iPSC characterization for donor pair 2.** iPSC characterization of *MAPT* p.R406W donor line (iPSC iPSC FA14-530.1.NGN2Δ1C05) and isogenic control (iPSC FA14-530.1Δ3G12.NGN2Δ1E04). Each iPSC line used in the study was verified by Sanger sequencing. Karyotyping was performed to confirm chromosomal stability. Immunocytochemistry was performed to confirm expression of pluripotency markers as well as NGN2 plasmid sequencing for integration confirmation.

**Supplemental Figure 3: Neuronal characterization.** iPSC-derived NGN2 neurons (DIV7) used throughout the experiments were characterized for neuronal markers, NGN2 neurons are MAP2-positive, Tuj1-positive and synapsin-positive by DIV14, detected by immunofluorescence staining.

**Supplemental Figure 4: LAMP1-positive vesicle characterization.** iPSC-derived NGN2 neurons (DIV14) used for detecting LAMP1+ vesicles by immunofluorescent staining using super resolution Airyscan microscopy. LAMP1+ vesicles (red) appear as donut-shaped structures in the soma as well as along the Tuj1+ neurites (green). LAMP1+ vesicles (green), showed to be mostly Cathepin-D positive (red), typically considered as lysosomes. CYTO-ID+ vesicles (green), are in close proximity to Lysotracker+ vesicles (red), zoomed inset of the merge image shows CYTO-ID+ (autophagosomes) and lysotracker+ (acidic vesicles) do not fully colocalize, therefor these structures are distinguishable (before and after fusion). Cells are ViaFluor^®^ 647 + (microtubule-magenta) neurons, DIV14, imaged live for CYTO-ID and Lysotracker, Airyscan super resolution microscopy, for the assay used in Figure 5 and 7.

**Supplemental Figure 5: list of genes differentially expressed in *MAPT* p.R406W neurons.** In *MAPT* p.R406W neurons, endolysosomal genes were enriched in pathways associated with phagosome maturation, regulation of autophagy, autophagosome and vesicle transport along the microtubule. List of genes coding for proteins in proton pump ATPases that were downregulated in *MAPT* p.R406W neurons. List of autophagy-related genes showed to be differentially expressed in *MAPT* p.R406W neurons.

**Supplemental Figure 6: G2-567 characterization in NGN2-neurons.** Neurons were treated with DMSO, G2-567 (0.5 µM for 14 days) or Bafilomycin (100nM for 2 hrs) as a control, fixed on DIV21. Neurons were immunostained for pTau levels (AT180-red) and MAP2 (green) and qualified for pTau intensity/cell (right). Neurons were also immunostained for pTau (AT180-green) inside LAMP1+ vesicles (red), and quantified for pTau intensity/lysosome upon treatment with DMSO, G2-567 and Bafilomycin. Lysosome size was also quantified in treated neurons by measuring LAMP1+ vesicle size (µm^2^). Kruskal-Wallis test followed by Dunn’s test. *, *p<0.05,* **, *p<0.001*, *\*\*\*\*, p<0.0001*.

**Supplemental Figure 7: Autophagy enhancer G2-567 rescues tau and autophagy defects in *MAPT* p.R406W neurons in an independent donor line.** *MAPT* p.R406W and isogenic control neurons were treated with G2-567 (0.5µM) and DMSO for 14 days beginning on DIV7, fixed on D21. A. Neurons (MAP2, green) stained for pTau (AT180, red). Scale bar 15µm. B. Quantification of pTau mean intensity per cell. Data are represented as mean ± SEM. Kruskal-Wallis test followed by Dunn’s test. **, *p*<0.01. C. Neurons were immunostained with MAP2+ (magenta) and JIP3 (green). Scale bar 10µm. D. Quantification of JIP3 total mean intensity. ANOVA, ***, p<0.0001. E. *MAPT* p.R406W and isogenic control neurons from DMSO control and G2-567-treated cells were probed for autophagosomes using CYTO-ID (green) and MAP2 (magenta). Scale bar 10µm. F. Quantification of CYTO-ID mean density. Kruskal-Wallis test followed by Dunn’s test, *\*\*, p<0.005, ****, p<0.0001.* G. Neurons treated with G2-567 were immunostained for MAP2 (magenta) and p62 (green). Scale bar 10µm. H. Quantification of p62 mean intensity. Kruskal-Walli’s test. *, *p*<0.05, **<0.01, ***, *p*<0.0001. I. Neurons treated with G2-567 were immunostained for MAP2 (magenta) and LC3B (red). Scale bar 10µm. J. Quantification of LC3B mean intensity per cell. Kruskal-Wallis test followed by Dunn’s test, **, *p*<0.01, ****, *p*<0.0001. All data are mean ± SEM.Data are representative of three independent experiments.

## Notes

### Competing Interest Statement

The authors have declared no competing interest.

