## Supplemental Figures for "Autophagic enhancer rescues Tau accumulation in a stem cell model of frontotemporal dementia"

### Supplemental Figure 1

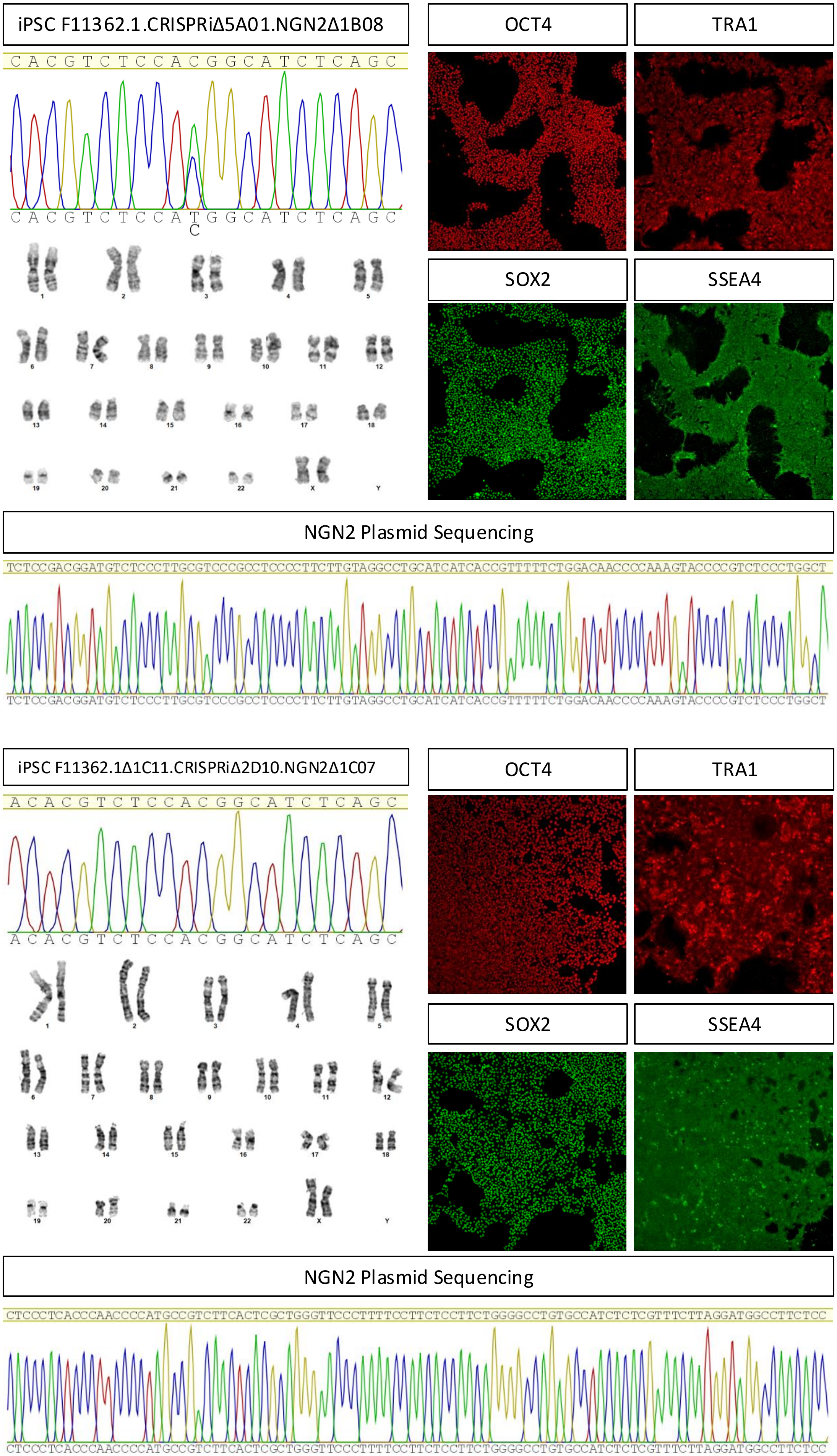

### Supplemental Figure 2

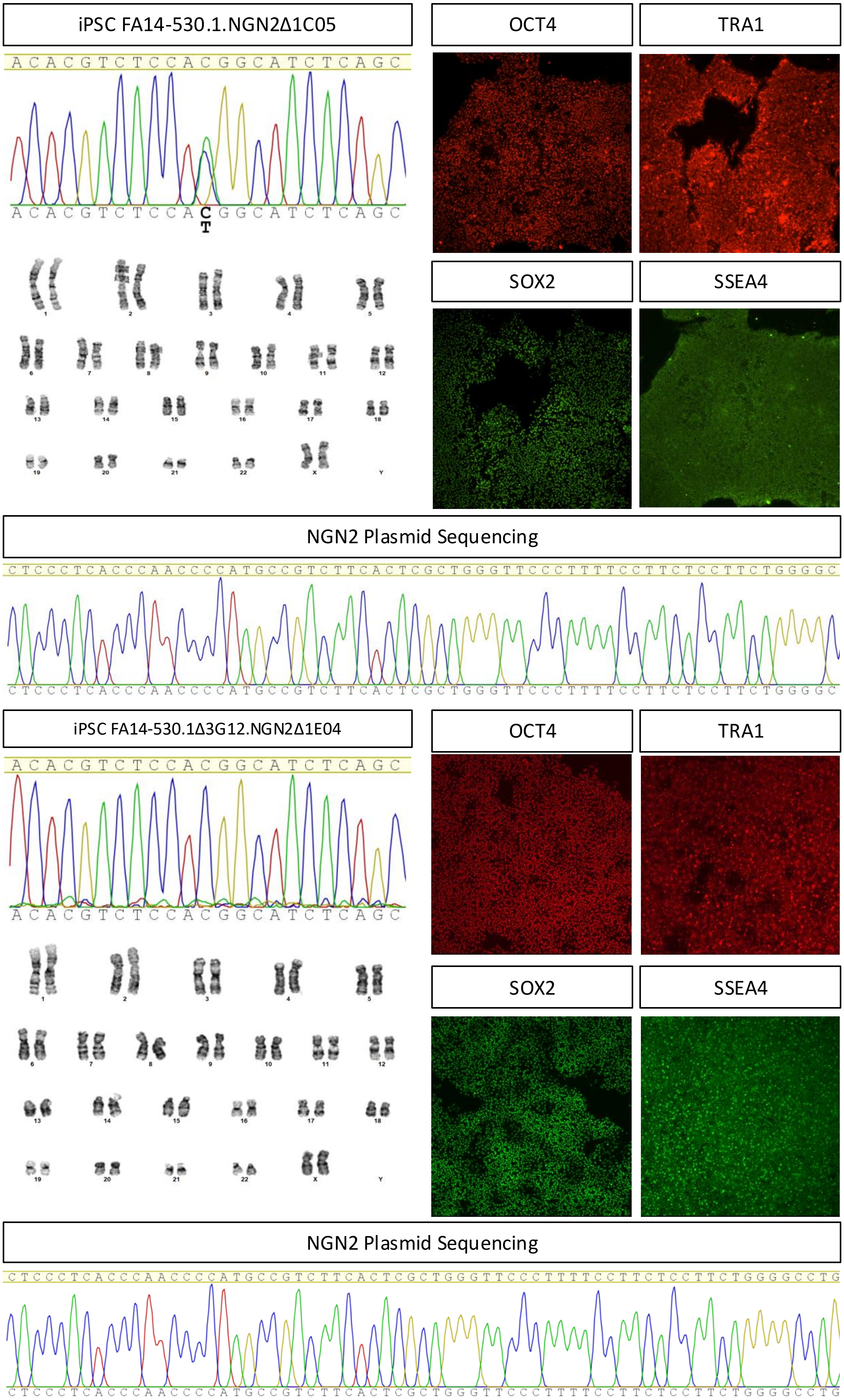

### Supplemental Figure 3

DIV14

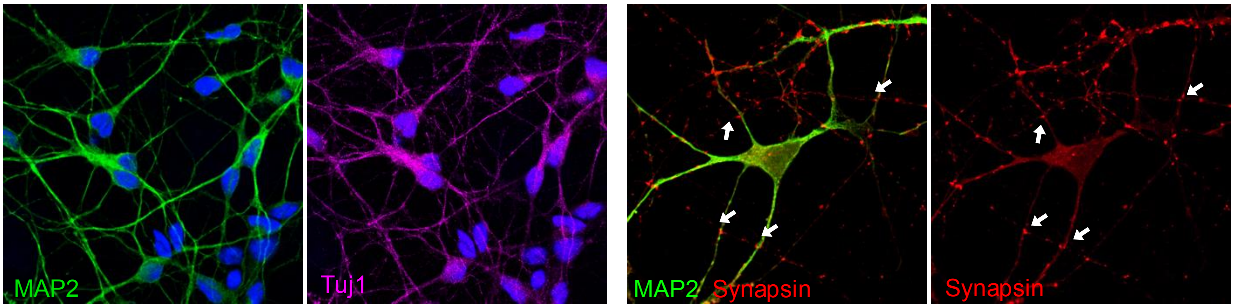

### Supplemental Figure 4

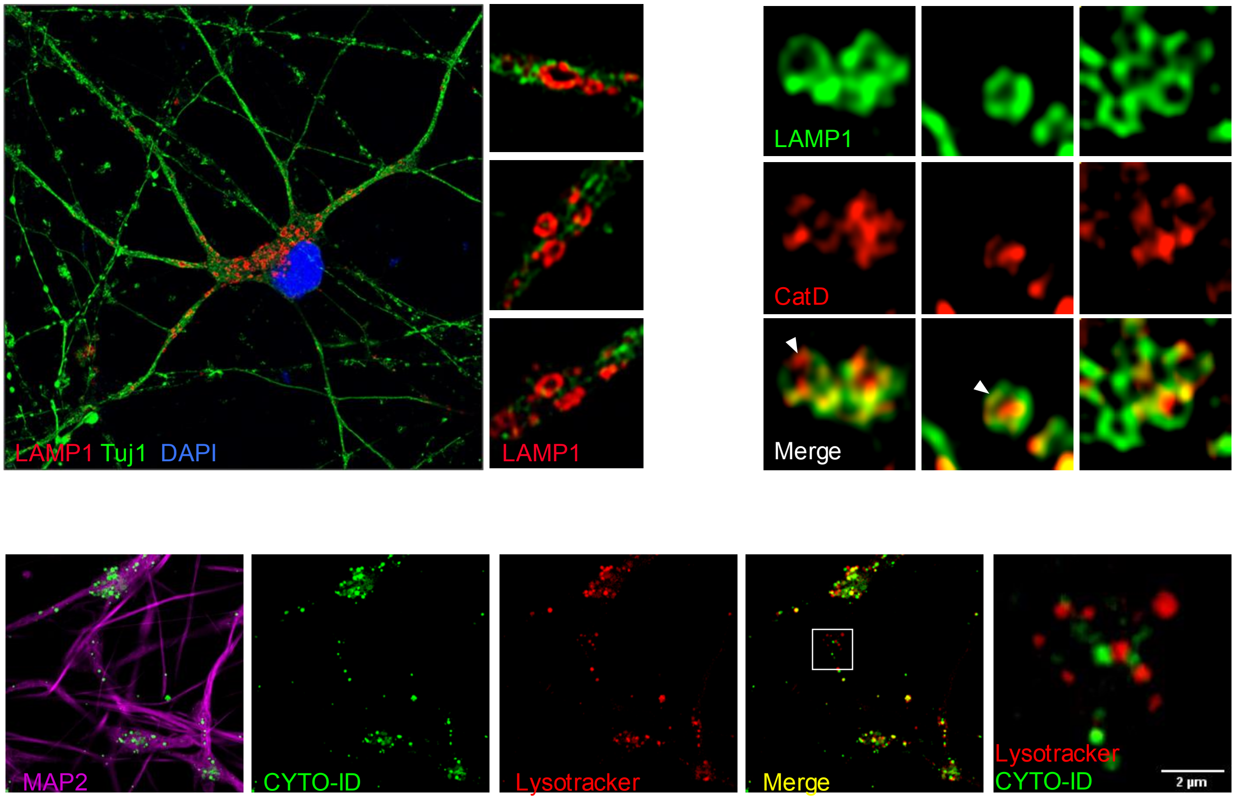

### Supplemental Figure 5

| Regulation | Analyses | Name/Description | p-value | Hit in Query List/userId |
|---|---|---|---|---|
| Up-Regulated | EnrichR | endonuclease activity | 2.044E-02 | CPSF4;XRCC2;DNASE1 |
| Up-Regulated | ToppGene | Golgi cis cisterna | 9.28E-05 | GOLGA8R,LLGL1,GOLGA8O,GOLGA8A |
| Down-Regulated | EnrichR | autophagosome | 5.327E-02 | VMP1;HSPA8 |
| Down-Regulated | EnrichR | lytic vacuole membrane | 4.686E-02 | HSPA8;ATP6V1B2;ATP6V1H;ATP6V1E1 |
| Down-Regulated | EnrichR | phagosome maturation | 8.341E-04 | ATP6V1B2;ATP6V1H;ATP6V1E1 |
| Down-Regulated | EnrichR | regulation of autophagy | 6.628E-03 | PLK2;ATP6V1B2;ATP6V1H;ATP6V1E1;EIF4 |
| Down-Regulated | EnrichR | vesicle transport along microtubule | 5.347E-03 | RAB1A;KIF3B |

#### genes coding for proteins in proton pump ATPase

| Regulation | Gene Name | log2 Fold change | p-value |
|---|---|---|---|
| Down-regulated | ATP10B | -0.45229009 | 3.196E-03 |
| Down-regulated | ATP1A3 | -0.253195718 | 3.647E-02 |
| Down-regulated | ATP5F1C | -0.18943298 | 5.206E-02 |
| Down-regulated | ATP5F1E | -0.19667578 | 3.616E-02 |
| Down-regulated | ATP5MD | -0.279564402 | 2.423E-02 |
| Down-regulated | ATP5PB | -0.251917081 | 8.919E-03 |
| Down-regulated | ATP6AP2 | -0.23102354 | 1.904E-02 |
| Down-regulated | ATP6V0A1 | -0.202284946 | 5.486E-02 |
| Down-regulated | ATP6V1B2 | -0.238001761 | 1.494E-02 |
| Down-regulated | ATP6V1C1 | -0.342021854 | 3.161E-03 |
| Down-regulated | ATP6V1E1 | -0.291136063 | 8.266E-03 |
| Down-regulated | ATP6V1H | -0.249047553 | 1.812E-02 |
| Down-regulated | ATP8A1 | -0.312282332 | 3.455E-03 |
| Down-regulated | ATP8B4 | -0.283422597 | 4.154E-02 |
| Down-regulated | ATP9A | -0.286120471 | 1.850E-03 |

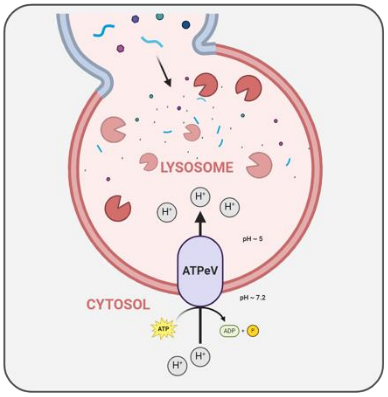


#### genes coding for proteins in Autophagy

| Regulation | Gene Name | log2 Fold change | p-value |
|---|---|---|---|
| Up-regulated | ATG16L2 | 0.54917053 | 2.324E-02 |
| Up-regulated | ATG2A | 0.396231508 | 1.087E-03 |
| Down-regulated | STX12 | -0.314246222 | 9.763E-03 |
| Down-regulated | STX7 | -0.310800269 | 2.016E-03 |
| Down-regulated | MAP1LC3A | -0.36849257 | 8.177E-03 |
| Down-regulated | SYT1 | -0.453961982 | 2.63E-06 |
| Down-regulated | SYT13 | -0.467925285 | 5.07E-05 |
| Down-regulated | SYT4 | -0.534948408 | 3.69E-07 |

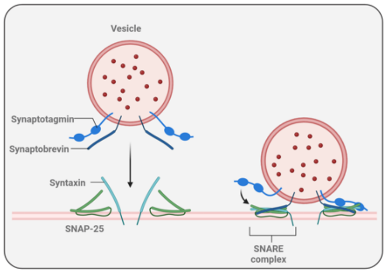

### Supplemental Figure 6

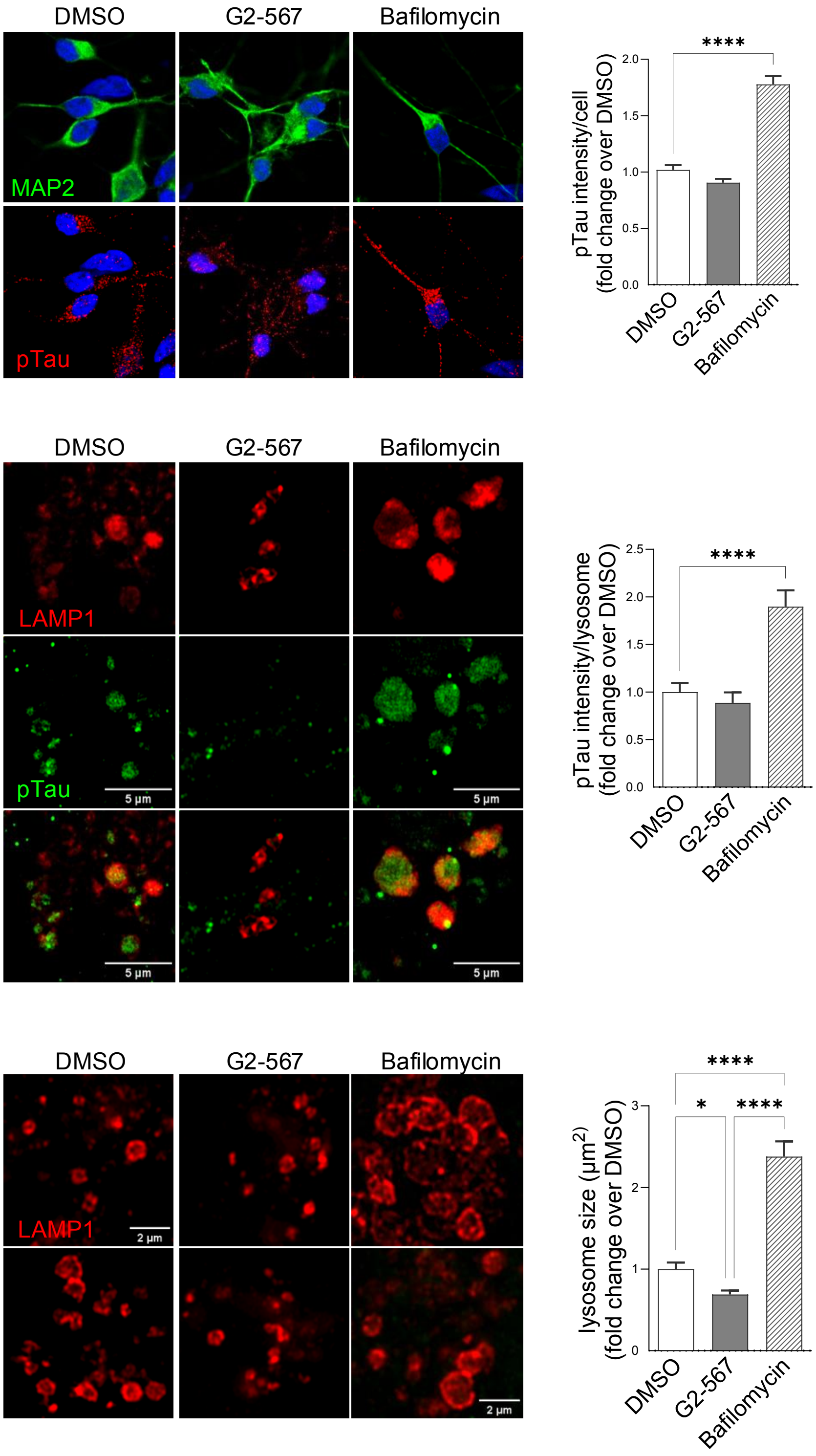

### Supplemental Figure 7

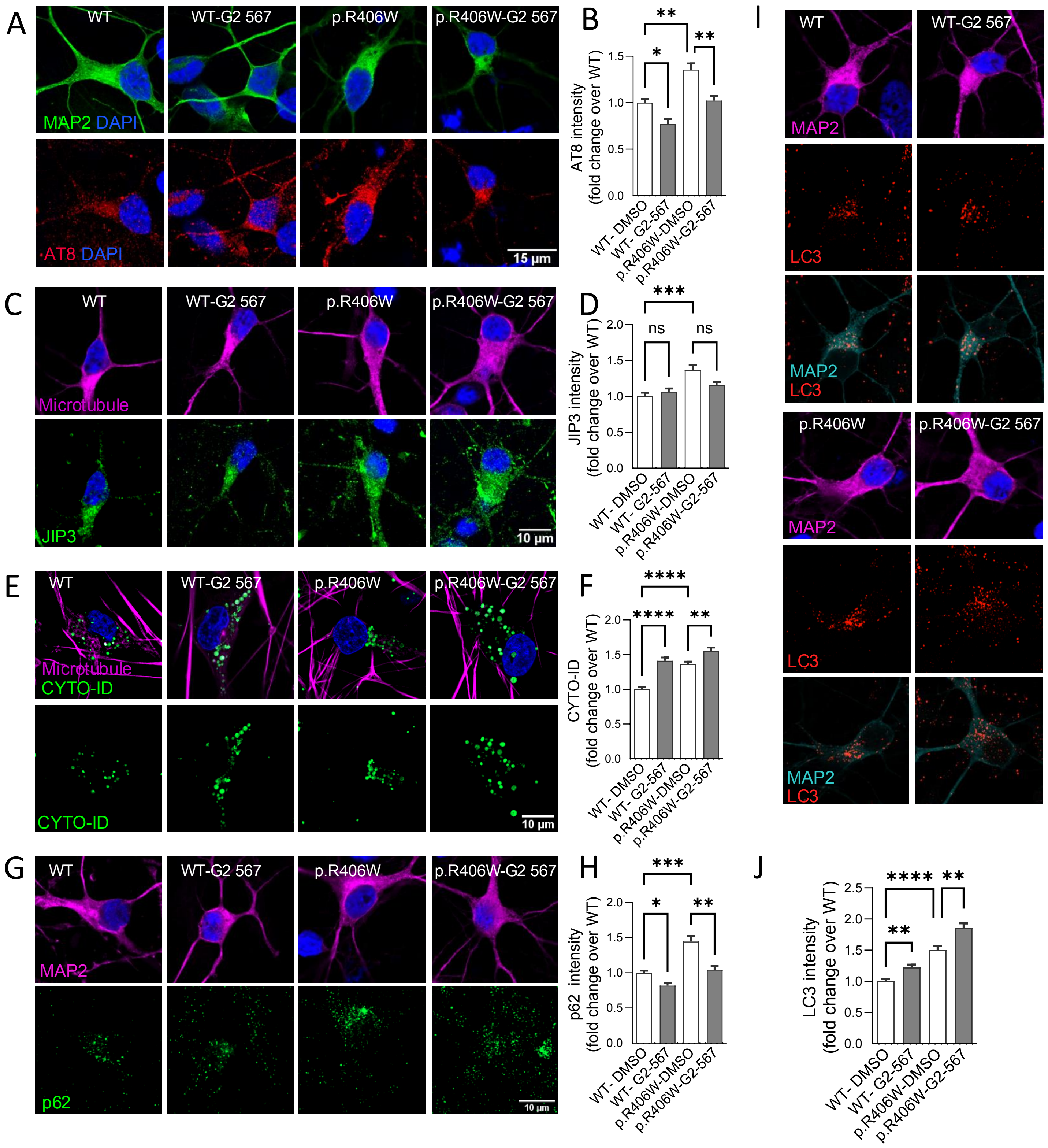
